# Direct Coupling Analysis and the Attention Mechanism

**DOI:** 10.1101/2024.02.06.579080

**Authors:** Francesco Caredda, Andrea Pagnani

## Abstract

Proteins are involved in nearly all cellular functions, encompassing roles in transport, signaling, enzymatic activity, and more. Their functionalities crucially depend on their complex three-dimensional arrangement. For this reason, being able to predict their structure from the amino acid sequence has been and still is a phenomenal computational challenge that the introduction of *AlphaFold* solved with unprecedented accuracy. However, the inherent complexity of *AlphaFold*’s architectures makes it challenging to understand the rules that ultimately shape the protein’s predicted structure. This study investigates a single-layer unsupervised model based on the attention mechanism. More precisely, we explore a Direct Coupling Analysis (DCA) method that mimics the attention mechanism of several popular *Transformer* architectures, such as *AlphaFold* itself. The model’s parameters, notably fewer than those in standard DCA-based algorithms, can be directly used for extracting structural determinants such as the contact map of the protein family under study. Additionally, the functional form of the energy function of the model enables us to deploy a multi-family learning strategy, allowing us to effectively integrate information across multiple protein families, whereas standard DCA algorithms are typically limited to single protein families. Finally, we implemented a generative version of the model using an autoregressive architecture, capable of efficiently generating new proteins in silico. The effectiveness of our Attention-Based DCA architecture is evaluated using different families of evolutionary-related proteins, whose structural data is sourced from the Pfam database.

## 1 Introduction

Proteins constitute a diverse category of biological compounds constructed from a set of 20 amino acids. Within an organism, they serve various functions, including structural support, mobility, and enzymatic activities. The effectiveness of a protein is intricately linked to its three-dimensional arrangement, known as its tertiary structure. This structure dictates the protein’s biological functionality when isolated and its interactions with other molecules within the cellular environment. As a consequence of the multitude of tasks proteins can perform, they lack a standardized three-dimensional conformation. Nonetheless, under physiological conditions, a protein’s three-dimensional configuration is influenced by its amino acid sequence [1]. Determining this dependence is theoretically and computationally challenging due to the system’s complexity. Only recently, the problem of predicting the folding of an amino acid sequence has seen a historical computational breakthrough thanks to *AlphaFold* in 2021 [2].

*AlphaFold* exploited decades of research in computational biology together with recent developments in machine learning. The most fundamental idea, which has been the center of this research area for years, is that evolutionary information can be extracted and used to determine patterns in phylogenetically related protein sequences (viz. homologs). Indeed, in the course of evolution, to preserve the functionality of a class of proteins, their structure must be conserved. Natural selection imposes constraints on single active sites or multi-amino acid motifs which are fundamental for the correct sequence folding. This leads to the idea of conservation and co-evolution [3], [4]. Given a Multiple Sequence Alignment (MSA) [5], single- and pair-wise frequencies of amino acids along different positions are enough to extract summary statistics that can be used to determine structural information by inferring the parameters of a Potts model, in what is commonly known as Direct Coupling Analysis (DCA) [6], [7]. The inference has been implemented in various ways and with various degrees of approximation during the last decade, leading to algorithms capable of determining the contact map of a protein family (PlmDCA, [8]), but also generating in silico sequences representative of the full statistics of the original protein family (bmDCA [9, 10], ArDCA, [11]).

A similar strategy is adopted by *AlphaFold* [2], where the self-attention mechanism allows for a direct representation of correlations over the MSA, albeit through multiple layers, [12, 13]. The attention mechanism was originally introduced in the context of Natural Language Processing (NLP) to overcome the limitations of sequential encoder-decoder architectures [14]. The basic idea is that long-range correlations within a dataset can be captured by a so-called *attention map*, encoding a custom functional relation between features of the dataset. For instance, in NLP the correlations to capture emerge at the semantic level [14], whereas in the case of structural protein inference from homology data, at the level of the individual residues as in the *Evoformer* architectural block in *AlphaFold*, in which a contact representation of the residues in a sequence is updated by conservation and co-evolution information extracted from an MSA and processed through self-attention layers.

## 2 Background and aims

In this article, we analyze the *factored attention layer* defined by Bhattacharya et al. [15] as a simplified version of the dot product self-attention mechanism [12]. In a factored attention layer, the positional degrees of freedom of the amino acid sequence are decoupled from the *color/amino acid* degrees of freedom representing each possible amino acid. This factorization separates the signal from a specific protein family and the signal due to the nature of amino acid interaction shared across multiple families. In their work, they demonstrate that a factored attention model can be traced back to a generalized Potts model [16], and ultimately all the methods developed in the context of DCA [7] can be used to obtain a contact prediction algorithm.

Interestingly, in [15] is shown that even if the number of parameters of the *factored attention* is significantly lower compared to the standard DCA models, their performance is almost equivalent. However, they limit their analysis to a contact score obtained in the same manner as for a generic Potts model, while here we push forward this analysis by studying the contact prediction obtained directly from the attention matrices of a factored attention model. More precisely, we show that the accuracy of the Frobenious and Attention scores is compatible across multiple protein families. We argue that this is a more direct way to understand the inner workings of more complex attention models, such as those used by *AlphaFold*. Moreover, we analyze the structure of the attention matrices to show their sparse nature in determining the structure of the protein family.

Another significant result found in [15] is that a factored model can be used to integrate signals from different protein families that can therefore share parameters. In particular, they showed that a set of shareable parameters can be learned from a protein family and then used with multiple other families for contact prediction, without loss in accuracy. Inspired by this, we introduce a multi-family learning scheme in which the shareable parameters are learned simultaneously on different protein families and used for contact prediction. This application exploits the factored nature of the model, highlighting the fundamental assumption that the signal from a protein family can be divided into two contributions: a family-specific signal and a universal, shareable signal that arises from structures and interactions common to all protein families.

In addition, we investigated the model’s hyper-parameter space, finding that it is characterized by an iso-performance phase diagram defined by the overall number of parameters. More precisely, by defining *H* as the number of heads and *d* as the inner dimension of the factored attention model, we observed that on the *H, d* curve induced by fixing the number of parameters, the contact prediction’s accuracy of the model is roughly constant (and improves upon increasing the total number of parameters, up to a certain threshold).

Finally, we introduced a generative version of the factored attention model by defining an autoregressive masking scheme inspired by the work in [11] where the same is applied to standard DCA. Sampling from the learned distribution produces generated MSAs that reproduce the same statistics as the natural ones.

As shown in Section 4, we argue that the attention mechanism and the Potts model are practically equivalent, in agreement with the recent theoretical evaluation using the Replica Method by Rende et al. [17]. Moreover, when modeling homology data, the transformer architecture gives possible advantages for analyzing the universal biochemical information which, if fully characterized and defined in terms of value matrices, could aid deep-learning machines such as *AlphaFold* itself.

Throughout the manuscript, we will use a Julia [18] in-house implementation of the factored self-attention mechanism that we refer to as *AttentionDCA*, to align this work to the terminology used in the field of DCA methods.

## 3 Methods

### 3.1 Factored Attention

As in every Direct Coupling Analysis implementation, sequences in a protein family are represented by a Multiple Sequence Alignment and can be thought of as independent samples (cf. Supplementary Material for details) from a probability distribution that we model as a Gibbs-Boltzmann measure over a Hamiltonian function defined as a Potts model [7]:

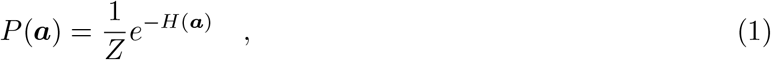

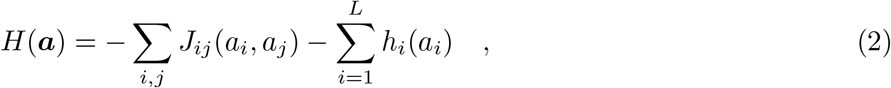

where, **a** = (*a*_1_, …, *a*_*L*_) is a sequence of *L* amino acids (*a*_*i*_ take value in an alphabet of 21 letters), J, and h are respectively the direct interaction tensor and the local field terms, while *Z* is a normalization constant, also known as partition function in the Statistical Physics jargon, given by:

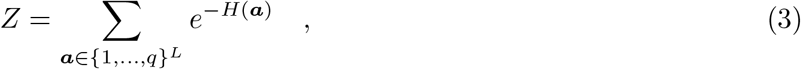

where *q* = 21 is the length of the amino acid dictionary, corresponding to the 20 natural amino acids plus a gap sign used during the alignment procedure in building the MSA. The inverse temperature *β*, which usually multiplies the energy term, is set equal to one, which is equivalent to implying its dependence directly inside terms *J* and *h*.

At this generic stage, tensor *J* ∈ ℝ^*L,L,q,q*^ encodes both positional and amino acid information, while *h* ∈ ℝ^*L,q*^ represents the local biases of each position in the sequence. To implement the factored attention mechanism discussed in [15], we discard the local field term, which is possible due to a gauge invariance of the parameters, and we write the interaction tensor mimicking the popular transformer [12] implementation of the attention mechanism for which:

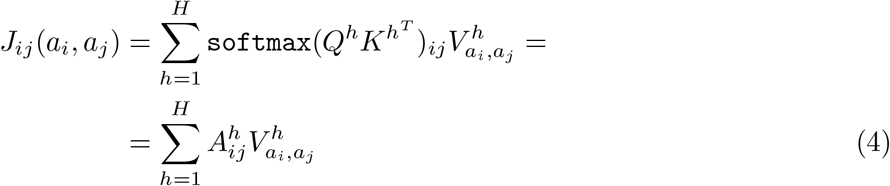

where, using the jargon from NLP, *Q*^*h*^ ∈ ℝ^*L,d*^, *K*^*h*^ ∈ ℝ^*L,d*^, *V* ^*h*^ ∈ ℝ^*q,q*^ are respectively the *query, key* and *value* matrices in one of *H* attention heads that build the interaction tensor *J*. The softmax is performed column-wise and the resulting matrix 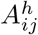, *i, j* ∈ 1, …, *L* (such that for all *i* ∈ 1, …, *L*,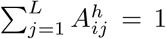) represents the self-attention of each pair of residues. The attention matrix in this form is meant to highlight the co-evolution relationships between different positions in a sequence, meanwhile a *M × M* column-attention would concern phylogenetic relations between different sequences in the same MSA; more on this can be found in Rao et al. [19] and Sgarbossa at al. [20].

A major implication of the decomposition of the interaction tensor into matrices *Q, K*, and *V* is that we are factoring the positional information contained inside matrices *Q* and *K*, depending on the specific protein family at hand, and the information contained inside matrix *V* which should capture the universal traits characterizing the interactions between the twenty natural amino acids. This turns out to be an interesting point for future development that plays an already crucial role in the multi-family version of the model discussed in Section 3.2.

To infer the parameters, i.e. train the model, we implement a pseudo-likelihood approximation, that can be interpreted as a precursor of the more general Masked Language Modeling scheme currently used for training state-of-the-art Large Language Models [21]. The full probability distribution is given by a factorization into single-site distributions conditioned to all the other sites in the sequence. The pseudo-likelihood of the model over a Multiple Sequence Alignment (MSA) 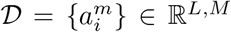 with *M* sequences (depth of the MSA) each of *L* amino acids (length of the MSA) can be written as

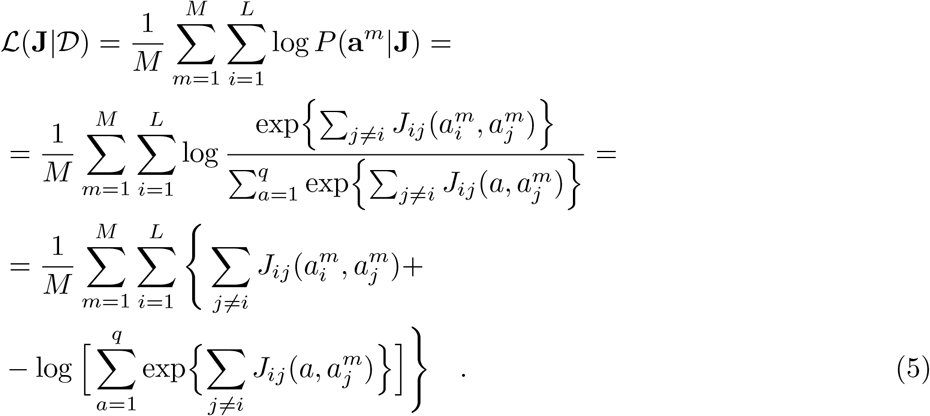

Finally, following a general procedure and in agreement with what is used in PlmDCA [8], we add an L2-regularization of the interaction tensor after an analysis of different regularization schemes. This penalizes large values of the *J*_*ij*_ matrix to avoid overfitting the direct coupling score, as defined in Equation 7. Thus, the total final likelihood is given by:

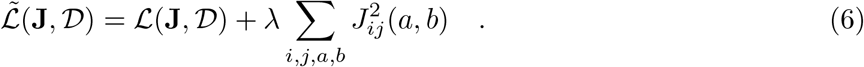

The decomposition of the interaction tensor given by Equation 4 has the side effect of making the total log-likelihood a non-convex-up function of its parameters, mainly due to the matrix products and the `softmax` function. Compared to models such as PlmDCA or ArDCA, this makes the maximization procedure much more challenging. To smooth out the complex landscape of peaks and troughs of the total likelihood during the maximization procedure, we implement an ADAM stochastic gradient ascent over mini-batches of fixed size [22].

Once the optimization is concluded and the parameters have been inferred, the standard way to obtain a contact prediction is by computing the *average-product-corrected* (cf. Supplementary Material) Frobenious norm of the interaction tensor which is interpreted as a score of the direct interaction between any position pair (*i, j*) in the chain [23]:

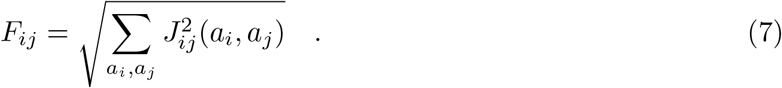

The higher the Frobenious score, the more likely two pairs are to be an actual contact in the structure. This is indeed what is shown in [15]. However, as mentioned in the introduction, we implement an alternative scheme to compute a contact score in terms of the positional-dependent attention matrices. This attention score can be defined as

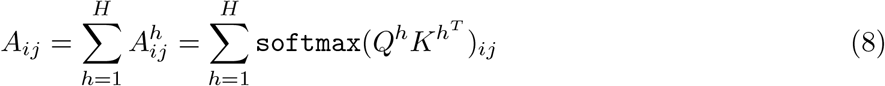

and it can be used in place of the original Frobenius score.

When testing the model on a protein family whose structure is known, we define the Positive Predicted Value (PPV) as the percentage of true positive (TP) contacts among the predicted ones: PPV(*n*) = TP(*n*)*/n*. The curve given by this measure, a function of *n*, represents the global accuracy of the model. Throughout the manuscript, we will use the consolidated notation PPV@*n* to indicate PPV(*n*).

Finally, since we are dealing with a heavily non-convex optimization problem, the results for different runs are generally affected by intrinsic noise. To reduce this, for each MSA we run the model several times and we build a final interaction score that represents a merging between the scores of each run. This is done by picking for each position pair its maximum score among all the runs. In Section 4 we compare PPV curves from single runs and merged multiple runs of AttentionDCA.

### 3.2 Multi-Family Learning

A long-standing goal of protein design is that of being able to selectively pick features from different protein families and generate sequences at first in silico and then in vivo so that these new artificial proteins reproduce those specifically chosen traits and functions [24], [25]. The first step toward this feat is that of designing a DCA model that can learn simultaneously from different MSAs. Standard DCA is not prone to this possibility since there is no obvious way to determine a set of parameters that can be shared among different families since both the interaction tensor and the local fields are family-dependent. However, in the Factored Attention implementation of DCA, the Value matrix

*V* ∈ ℝ^*q,q*^ does not depend on the specific family. In [15] it is shown that a set of value matrices learned in a specific protein family can be *frozen* and used during the learning of Query and Key matrices for another protein family. Motivated by this we introduce a model in which the Value matrices are shared and learned across different MSAs simultaneously. Given *N*_*F*_ protein families, a simple implementation of this is obtained by defining a multi-family likelihood ℒ_*MF*_ given by the sum of the *N*_*F*_ single-family likelihoods coupled by the same set of Value matrices {*V* ^*h*^}_*h*=1, …,*H*_:

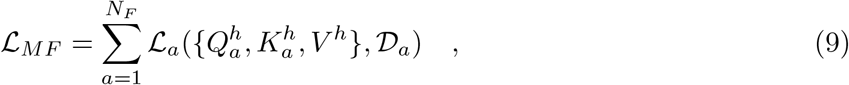

where each single-family likelihood is given by Equations 4 and 5. The set of parameters 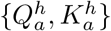, one for each family, and {*V* ^*h*^}, shared across all families, can be used to extract the contact score of each MSA used during the learning. The 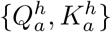 matrices must each have the same number of heads so that it is possible to share a common set of 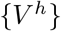 matrices among them. In Section 4 we show the results of this PPV compared to those obtained from standard single-family learning. Pseudo-likelihood maximization is used for inferring both the shared and family-specific parameters.

### 3.3 Generative model

The maximum pseudo-likelihood criterion used for the inference in PlmDCA and AttentionDCA does neither provide a fast way to sample the probability distribution nor an accurate sample due to the approximation used to factorize the probability distribution. Because of this, to develop the generative version of the model we started from ArDCA [11], which is so far the state of the art for generative DCA architectures. In particular, it exploits a simple autoregressive model enforced by the exact decomposition

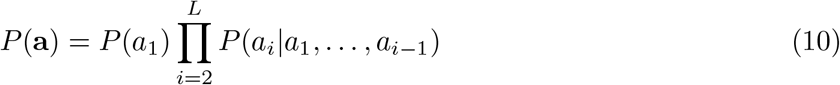

with

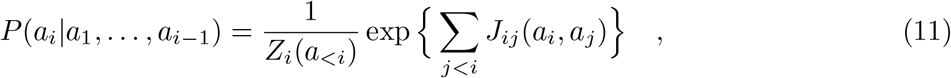

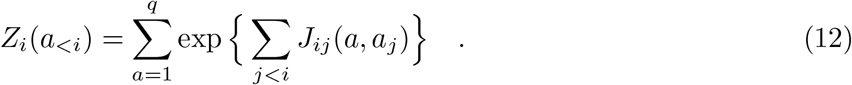

This form allows for fast computation of the single-site partition function and a quick autoregressive sampling from *L* single-valued distributions.

To exploit the already existing libraries of ArDCA and the architecture for AttentionDCA, we apply a multiplicative mask to the interaction tensor to implement an autoregressive structure of the distribution, i.e. so that *J*_*ij*_ is a lower-triangular matrix with zeros above and on the diagonal:

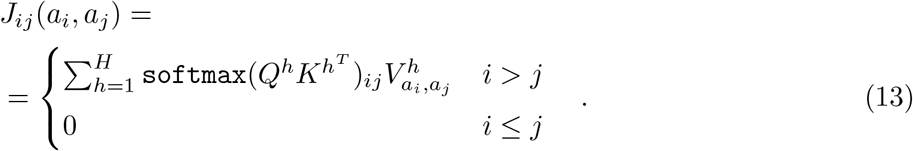

As discussed in Trinquier et al. [11], the resulting interaction tensor cannot be interpreted as a matrix of direct couplings in the same way as in standard DCA, mainly because in the autoregressive implementation, the interactions of each position in the chain are conditioned only to partial sequences instead of being conditioned to all other amino acid positions. Therefore, the common technique to predict the residue-residue contacts is to extract coupling information directly from the epistatic score evaluated from the model. For amino acids *b*_*i*_, *b*_*j*_ at positions (*i, j*), their epistatic score is defined as the difference between the effects of simultaneous mutations on both sites and the sum of the single site mutations when introduced in the wild-type **a** = (*a*_*i*_, …, *a*_*j*_) as in:

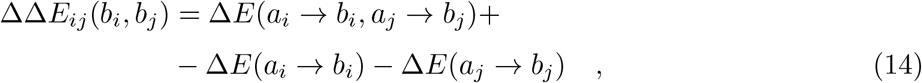

where the energy difference for a given mutation *a*_*i*_ → *b*_*i*_ is given by:

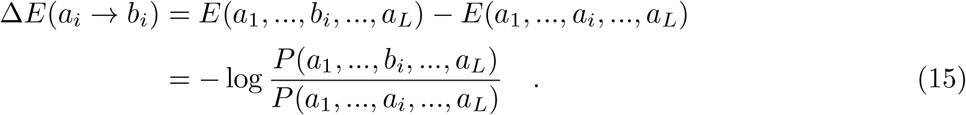

The ΔΔ*E*_*ij*_(*b*_*i*_, *b*_*j*_) replaces the interaction tensor *J*_*ij*_(*a*_*i*_,*a*_*j*_) in the Frobenious norm, cf. Eq. 7, used to build the Epistatic Score *ES*_*ij*_ so that a contact map can be produced and compared with the true structure of the protein. It is worth mentioning that the choice of the wild-type sequence in the definition of the Epistatic Score is practically immaterial: in Supplementary Material Sec. D we show that different choices of the wild-type reference sequence and even a randomly generated one produce equivalent results. Finally, an alternative to using the Epistatic Score to produce a contact score from the generative model is that of sampling an artificial MSA to be fed directly to a non-generative version, either PlmDCA or AttentionDCA, to define a meaningful Frobenious Score. Both alternatives will be discussed in the next section.

## 4 Results

The model, in its standard and autoregressive implementations, has been tested against nine protein families taken from the Protein Data Bank. Each family is represented by an MSA whose length *L* (number of amino acids per sequence) and depth *M* (number of sequences per MSA) vary, Table 1 summarizes some of the main information regarding each family. A crucial parameter is the effective depth *M*_*eff*_ that quantifies the effective number of independent sequences in the MSA. As we show in the following, the effective depth turns out to be crucial in determining the accuracy of the Epistatic Score contact prediction for the autoregressive generative version of the model, cf. 4.2.

**Table 1:**
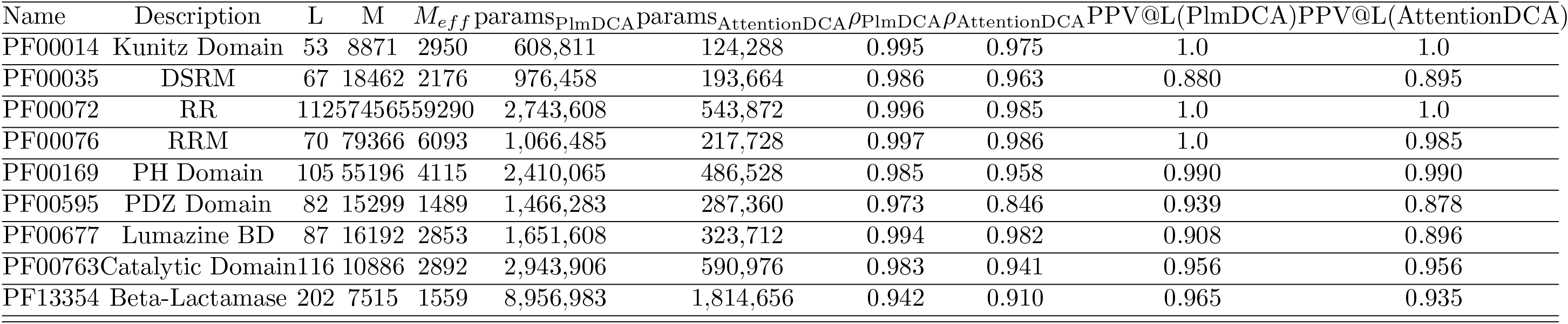
Summary of the protein families used throughout the analysis and results (MSA’s length, depth, effective depth, number of parameters in PlmDCA, number of parameters of AttentionDCA at *c*_*r*_ = 0.2, two-site connected correlation Pearson coefficient for ArDCA and AttentionDCA, Frobenious score positive predicted values at *L* for PlmDCA and AttentionDCA).

### 4.1 Standard version

#### 4.1.1 Parameter reduction and hyper-parameters

Concerning the hyper-parameters of the learning process, we use a learning rate *η* = 0.005 and a mini-batch size *n*_*b*_ = 1000, while the number of epochs varies depending on the depth of each MSA, cf. Supplementary Material. A more insightful analysis regards the hyper-parameters of the model itself, i.e. the number of heads *H* used to define the interaction tensor and the inner dimension *d* of the rectangular matrices *Q* and *K*. In standard DCA models, the size of the interaction tensor scales quadratically with the length of the protein family. However, empirically the number of contacts in a protein is proportional to its length *L* (and not to *L*^2^) [26], therefore we can expect the interaction tensor to be effectively sparse. In AttentionDCA, due to the low-rank decomposition of the interaction tensor, the scaling of the parameters is linear with the size of the family L: *N*_AttentionDCA_ = 2*HLd* + *Hq*^2^. The parameter compression ratio relative to PlmDCA can be computed by fixing the value of *d* and *H*:

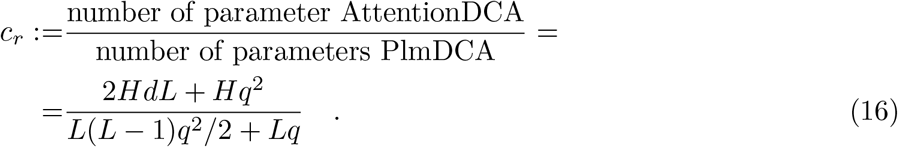

During the process of tuning the hyper-parameters of the model, we observe that, at a fixed compression ratio, the precision of the contact prediction is constant regardless of the choice of *H* and *d*. Therefore the number of parameters determines the quality of the model. Although a similar analysis regarding the dependence of the contact prediction’s accuracy on the number of heads of the model is present in [15], our findings show that it isn’t the number of heads that influences the accuracy of the model, but its global number of parameters. However, increasing the compression ratio, i.e., lowering the distance between PlmDCA and AttentionDCA in terms of the number of parameters, does not improve the contact prediction’s accuracy indefinitely. Indeed, having the same number of parameters of PlmDCA results in sensibly worse results. This may be because the likelihood in this architecture does not have a clear absolute maximum since the function is non-convex-up. Increasing the parameters enhances this condition making the inference harder and harder. After some trials, we conclude that the optimal parameter compression lies between 5% and 20%. Focusing on families PF00014, PF00076, and PF00763 respectively, Figure 1 shows the PPV evaluated at *L*, 3*L/*2, and 2*L* for different values of the head number *H* and the inner dimension *d* so that the parameter compression is fixed at values *c*_*r*_ = {0.05, 0.15, 0.25, 0.35}. It can be seen how a plateau is reached for *H >* 10, effectively highlighting the fact that performances remain constant at fixed compression, regardless of the hyper-parameters *H* and *d*. Another interesting feature is that for *H* ≲ 10, the value of the PPV is significantly lower than otherwise. This is due to a non-reliable inference of the parameters, as it can be seen that soon after the beginning of the gradient ascent, it hits a barrier and stops. The results shown in Figure 1 are averaged across multiple realizations, even though for most points the error bar is too small to be noticeable.

**Figure 1:**
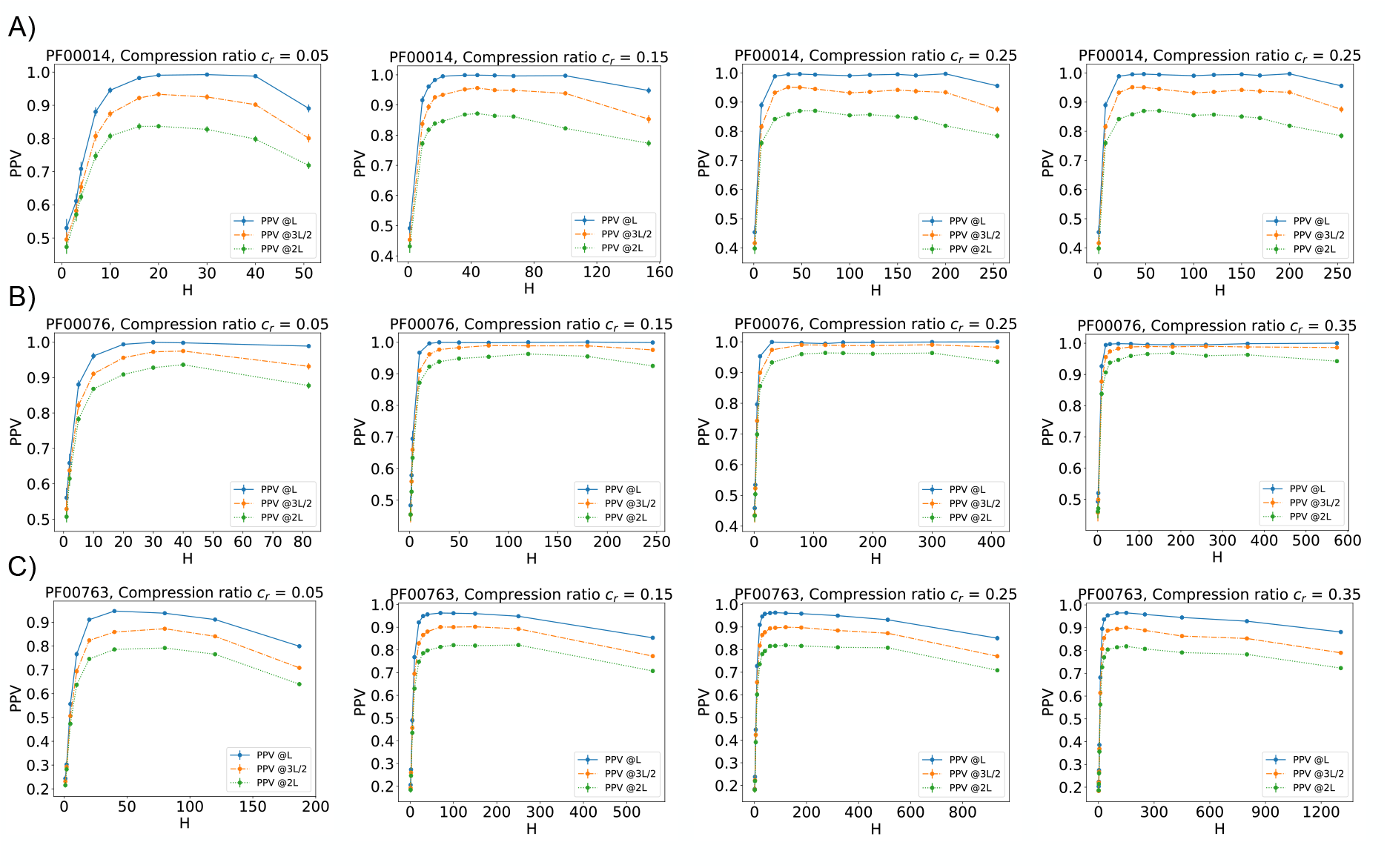
PPV @L, 3L/2 and 2L (blue, orange and green curve respectively) for different values of the compression ratio *r* for families PF00014, PF00076 and PF00763 respectively in panels A), B) and C). For each value of *H* there is a corresponding value of *d* given by Equation 16 at fixed *c*_*r*_.

#### 4.1.2 Contact Prediction

The standard way to perform a contact prediction for DCA methods is to compute the Frobenious norm of the inferred interaction tensor after a transformation which brings the parameters in zero-sum gauge, the gauge that minimizes the Frobenious norm, and a consecutive *average product correction* [8]. Sorting the Frobenious score (FS) results in a list of contacts that can be compared to the actual known structure of the protein family to produce a PPV curve. Alternatively, the factored attention form of the interaction tensor suggests another way of determining the contacts of a protein family. Averaging the positional-dependent attention matrix through each head results in a matrix of the form given by equation 8, the attention score (AS). After the *average product correction*, the contact prediction is given by the list of contacts (*i, j*) sorted by highest *A*_*ij*_. As already mentioned, the non-convexity of the problem and the roughness of the energy landscape produce a noise in the final inference which can be smoothed out by averaging through different realizations. In particular in the case of the FS contact prediction we compute the score for each realization of the inference and, for each position pair (*i, j*), we define its score as the maximum through all realizations: 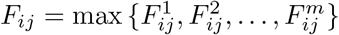 among the resulting ones, so that to produce a final merged scored which results in the most likely. In the case of the AS contact prediction, it is sufficient to simply average the resulting attention matrices of each realization. Following an analysis with an increasing number of realizations *m*, we identified an accuracy threshold at *m* = 20, which was subsequently used for the simulations.

Figure 2 shows the comparison between the PPV curves extracted from different implementations of AttentionDCA and those from PlmDCA (black curve), which serves as a benchmark model, applied to families PF00014, PF00072 and PF00763. Gray curves represent an ideal perfect model which predicts all and only positive contacts, while the red and blue curves are respectively the FS PPV from the merged- and single-realization of AttentionDCA. These results have been obtained by setting the compression ratio between PlmDCA and AttentionDCA to 20%, and for each family, apart from fluctuations in the single-realization curves, the results are in very strong agreement with PlmDCA, despite the significant parameter reduction between the two models. Dark and light green curves represent the PPVs evaluated from the multi-family leaning discussed in Section 3.2. In particular, the learning has been performed in parallel among all families using a fixed number of heads *H* = 128 and varying the inner dimension *d* to match the single-family learnings at 20% compression ratio. The shown curves are obtained through the Frobenious score computed on the *J*_*ij*_ tensor which is given by Equation 4 and parameters 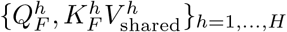, for *F* = {PF00014, PF00072, PF00763}.

**Figure 2:**
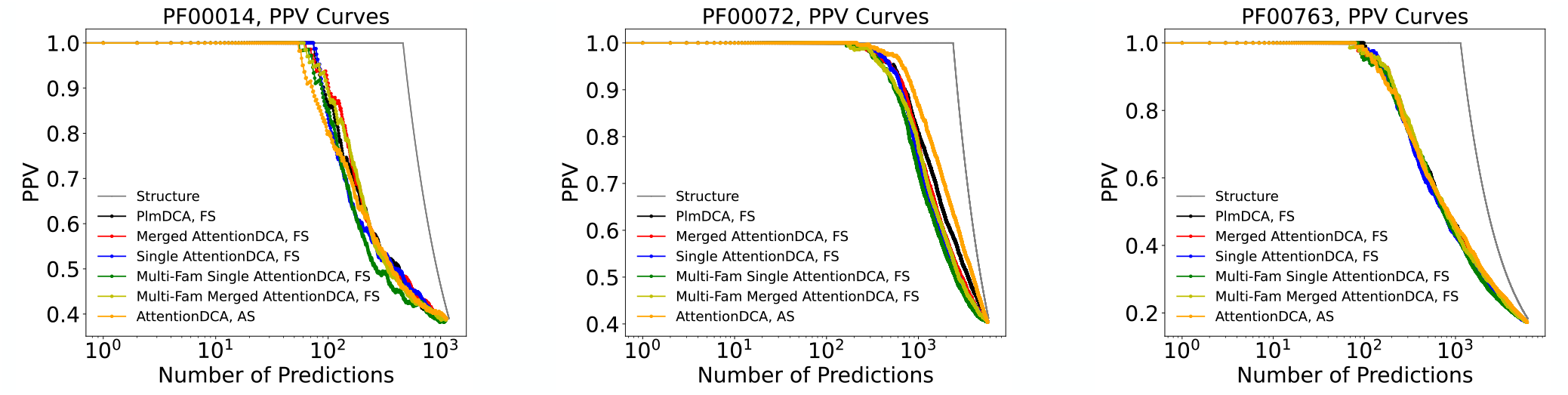
Positive Predicted Value Curves for families PF00014, PF00072, PF00763 respectively. Each plot compares PPVs computed through the Frobenious Score (FS) and the Attention Score (AS), in particular: PlmDCA with FS (black curve); merged and single AttentionDCA with FS (red and blue curves respectively); Multi-Family AttentionDCA with merged and single realization (green and yellow curves respectively); AttentionDCA with AS (orange curve). The gray curve represents the PPV of an ideal model that predicts all and only positive contacts, hence it depicts the prediction of the full structure of the family.

#### 4.1.3 Attention Heads

In the field of Natural Language Processing [27], the self-attention mechanism is used to learn custom functional relationships between the elements of a dataset to produce an encoded description of the input itself [28]. In the context of protein structure prediction, the attention mechanism used in *AlphaFold* untangles the physical and evolutionary constraints between phylogenetically related sequences [29, 30]. Since the factored attention model is effectively a single-layer self-attention architecture, we expect the attention heads inside the *J*_*ij*_ tensor to directly capture some level of structural information that could be compared to the actual contact map obtained by the average-product-corrected Frobenious norm of the interaction tensor. Ideally, one would expect each head to focus its attention on a specific spatial structure in the same way as a transformer head would specialize in different semantic functions when used in the context of NLP [31]. However, it is extremely difficult to determine if this is the case for protein attention due to the lack of an obvious interpretation key between the several possible structures arising in a protein and the variable number of heads. Also, given the form of Equation 4, a single head generally produces an asymmetric attention matrix, hence it is hard to understand which are the structures emerging from each one of them. A solution to both these problems is to simply average out all heads into a single attention matrix as in Equation 8.

The average attention matrix can be used to extract structural information by producing a list of contacts as discussed in subsection 4.1.2 and shown by the orange curves in Figure 2. However, the most direct way in which the average attention matrix can be interpreted is by comparing it to the actual contact map of the protein family. Figure 3 compares the contact map built from a significant fraction of predicted contacts and the heat map of the attention matrix for families PF00014, PF00072, and PF00763. In the contact maps, blue and red dots represent the positive and negative predictions, while gray dots show the actual structure of the family. Conversely, the heat maps are displayed on a grayscale, while the actual structure is given by the red dots underneath. It can be seen that the maxima of the attention matrix, i.e. the darker dots, lie within the structure of the protein, proving that the attention model is focused on the structural contacts. Even though, as already mentioned, the specialization of single attention heads is not trivial to understand, an important feature that can be observed is the sparsification of the actual structural information contained in each one of them. A quantitative analysis of the sparsity of the attention heads can be as follows: given {***A***^*h*^}_*h*=1,…,*H*_ (such that for each *h*, 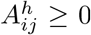, *i, j* = 1, …, *L*), for each head we select the *k* highest elements. This results in a list *S* of *k × H* elements that can be used to build a new attention matrix [*Ã*]_*i,j*=1,…,*L*_ either by maximizing or averaging for each position pair (*i, j*) over the subset of *S*_(*i,j*)_ ⊂ *S* which contains all the elements relative to that particular (*i, j*):

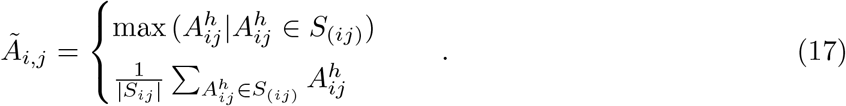

The resulting matrix is then *average-product* -corrected, symmetrized, and used as a contact score to produce a PPV curve and a contact map. By varying the value of *k*, i.e. the number of elements selected from each attention head, it is possible to evaluate the sparsity of the matrices by comparing the positive predicted values relative to each *k*. Figure 4 shows the PPV evaluated at *L* and 2*L* for increasing values of *k* for families PF00014, PF00072 and PF00763. Results are compared with the corresponding values from the standard Frobenious norm approach by AttentionDCA. It can be seen that when using the maximum to construct the sparse matrix, a very small fraction of the total elements is enough to return a PPV that is close to that obtained from the Frobenious norm. On the other hand, the PPV computed from the mean-version of the sparse matrix is slower to grow, but again for 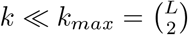 it reaches the accuracy of standard FS PPVs. This means that regardless of the version used for constructing *Ã*, a small number of elements is enough to capture structural information comparable to the full information obtained by the Frobenious norm. Figure 5 shows the full PPV curves from sparse attention matrices computed on a restricted number of significant values of *k*: 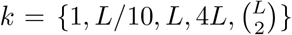. Furthermore, Figure 3C) shows the heatmap of the sparse attention matrix constructed using the maximum version with *k* = *L*.

**Figure 3:**
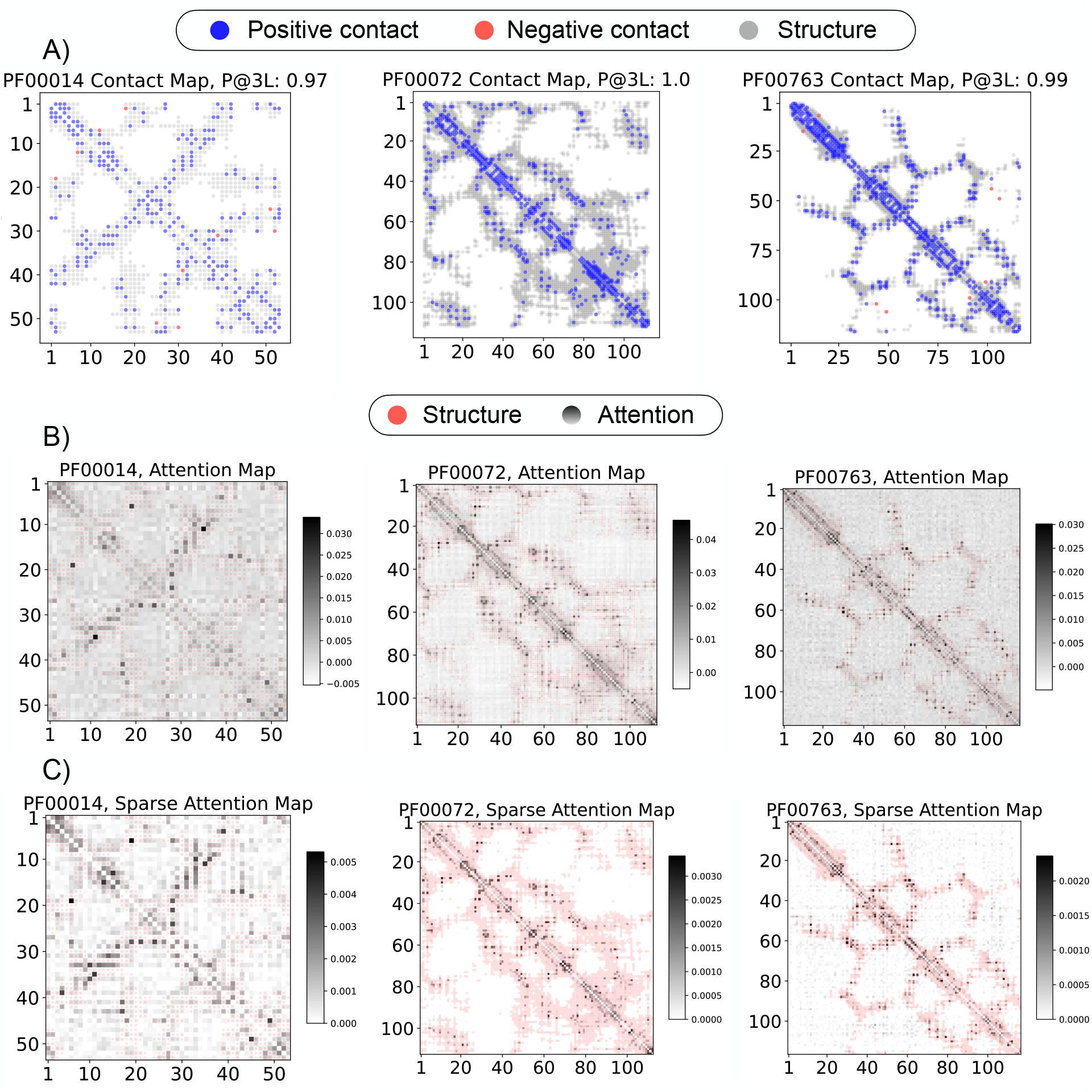
Various contact maps for families PF00014, PF00072, PF00763. A) Contact map built from the Frobenious Score applied to the interaction tensor by AttentionDCA. Blue and red dots represent positive and negative predictions respectively, while gray dots reproduce the structure of the family. B) Grayscale heat map of the averaged attention matrix built using Equation 8, from the {*Q*^*h*^, *K*^*h*^} _*h*=1,…,*H*_ matrices by AttentionDCA. C) Grayscale heat map for the sparse attention matrix constructed by maximizing over the *k* highest elements from each head, using *k* = *L*, as in Equation 17. Both in B) and C) structural information is depicted in red.

**Figure 4:**
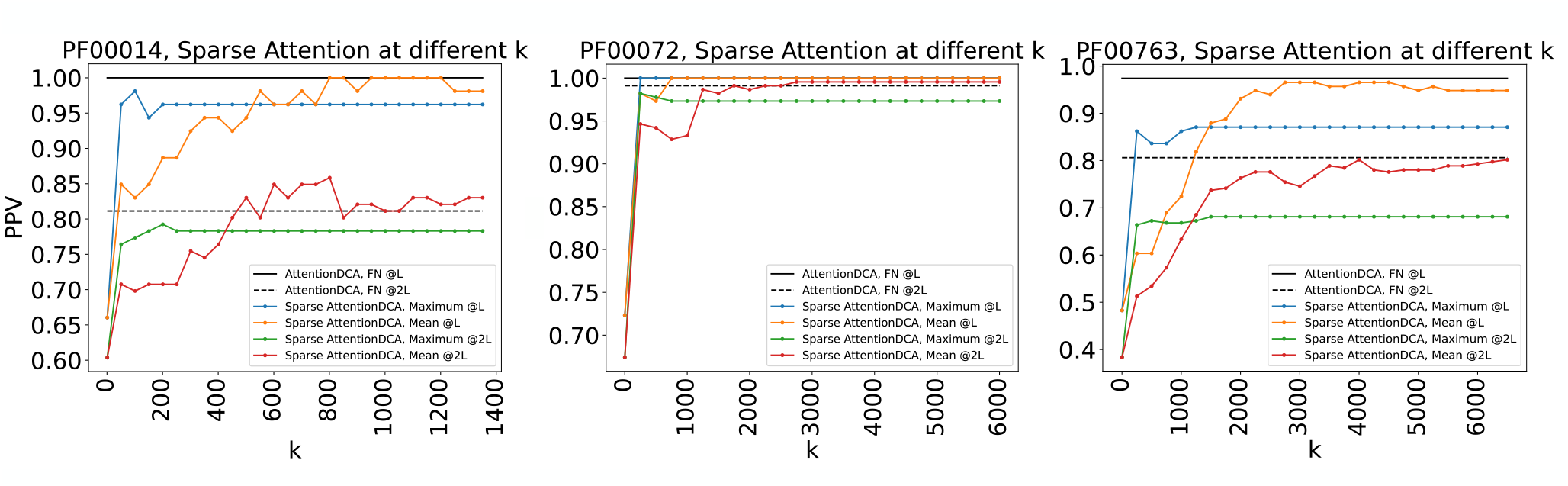
PPV curves @L and 2L for the average- and maximum-sparse Attention Score as the value of the number of elements k from each varies. Blue and orange curves represent the PPV @L for the average- and maximum-sparse version respectively and are to be compared to the full black line representing standard AttentionDCA computed using the Frobenious Score (FS). Similarly, green and red curves represent the PPV @2L and are to be compared to the dotted black line. Families PF00014, PF00072 and PF00763 are shown.

**Figure 5:**
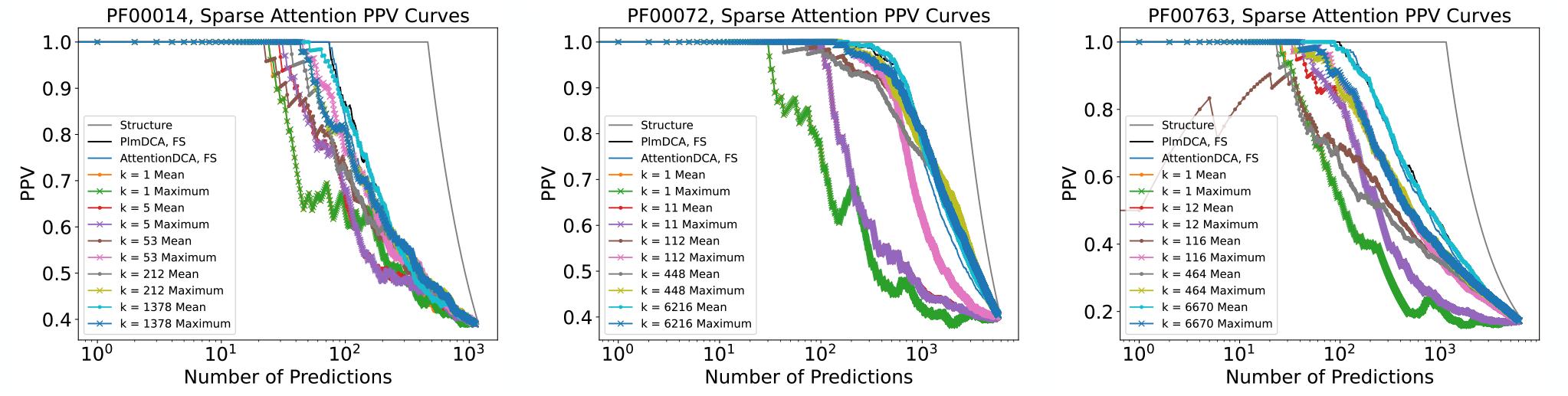
PPV curves of families PF00014, PF00072, PF00763 for the average- and maximum-sparse Attention Score computed for 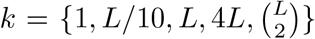 and compared to the standard Frobenious Score (FS) from PlmDCA and AttentionDCA (black and blue curves respectively).

### 4.2 Autoregressive generative version

Even though the autoregressive version of our architecture can be used for contact prediction, its main target is to efficiently sample the distribution to generate large MSAs matching the summary statistics of the natural MSAs used for inference.

As described in Section 3.3, the form of the interaction term *J*_*ij*_(*a, b*) of the autoregressive version does not allow for a meaningful mapping to direct interactions among amino acids. Therefore, the epistatic score measure defined in Equation 14 must be used to define a list of predicted contacts and the corresponding PPV curve. Figure 8A) shows different PPV curves computed on families PF00014, PF00072 and PF00763. In particular, it compares the performance of PlmDCA (black curve) and ArDCA (red curve), both acting as benchmarks, and those of the autoregressive AttentionDCA in the single- and merged-realization implementations (blue and green curves respectively). Along with the epistatic measure, another way to perform a contact prediction is to produce a generated MSA, sampled using AttentionDCA, and feed it to PlmDCA or AttentionDCA itself (in its non-generative version). This last possibility is represented by the yellow curve and it is among the best options for contact prediction within the autoregressive versions, either AttentionDCA or ArDCA. This can also be thought of as a way to test the accuracy of the generative capabilities of the autoregressive model, which exploits the exact decomposition in Equation 10 to sample from *L* single-valued probability distributions. Moreover, the generativity can be assessed by comparing the summary statistics of natural and generated MSAs. The most informative summary statistics are given by the connected two-site correlations of amino acids in different positions of an MSA, cf. Supplementary Material. Their comparison is shown in Figure 8B), along with their Pearson’s correlation coefficient, proving the accuracy of the model. Finally, a PCA analysis can be performed to show that the principal components of the generated data are comparable to those of the natural data. Figure 6 shows the comparison between the two principal components of the natural data and the artificial data generated by ArDCA, used as a benchmark model, and by AttentionDCA on families PF00014, PF00763 and PF13354 (cf. Supplementary Material for the other families).

**Figure 6:**
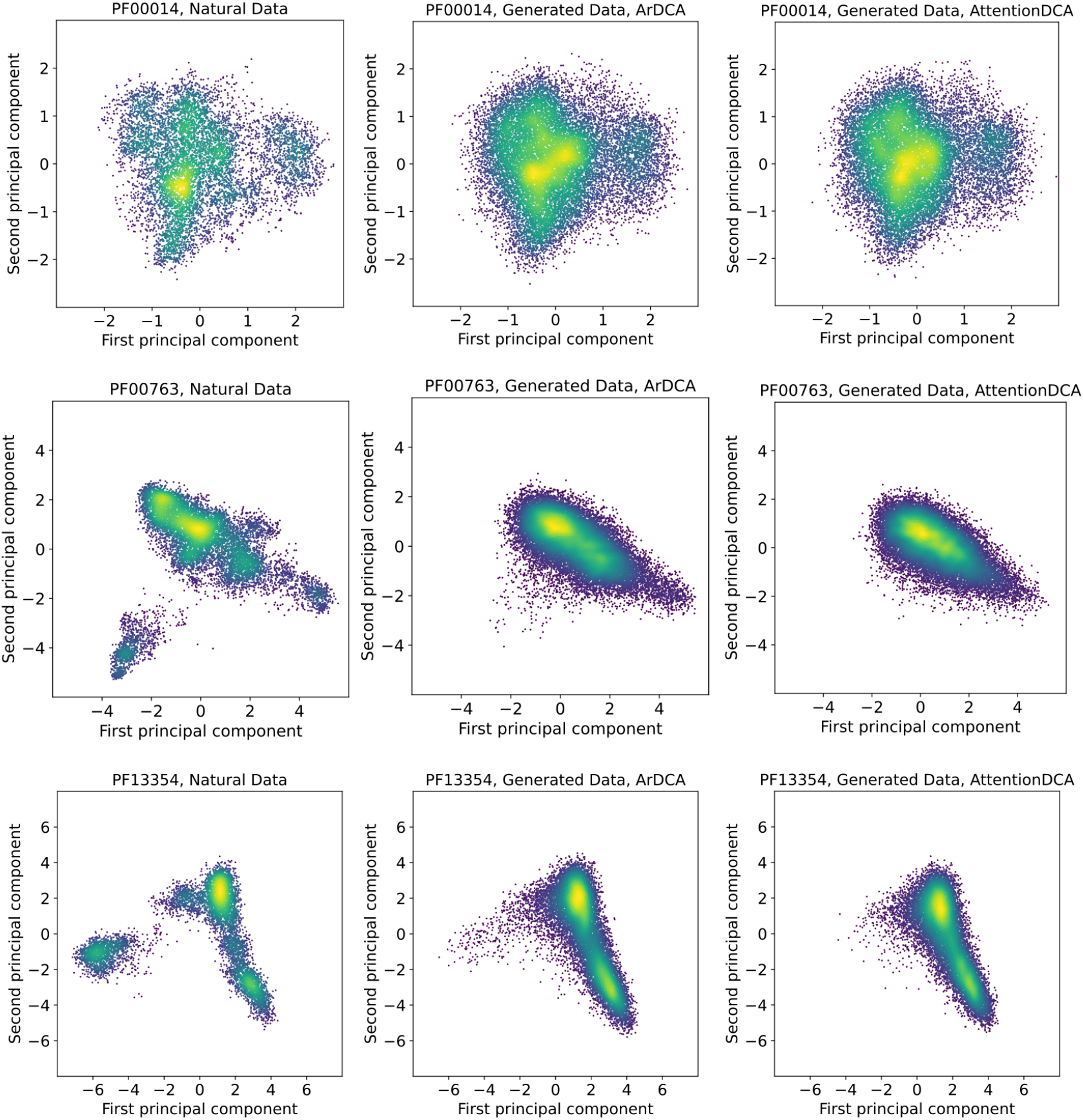
Principal Component Analysis on protein families PF00014, PF00763 and PF13354. Comparison between natural data and artificial data generated by using ArDCA and AttentionDCA.

For many families among the ones described in Table 1, the results of the contact prediction from the epistatic score lie significantly below those from the Frobenious score of the standard version of AttentionDCA or PlmDCA. The reasons for this can be found in the dependency of the autoregressive version on the *M*_*eff*_ of the family. If one were to reduce the effective depth of an MSA, after a certain threshold, the DCA inference would suffer a lack of observations and the contact prediction would get less and less reliable. Different DCA models and different inference techniques have variable *M*_*eff*_ thresholds. Figure 7 shows how the accuracy of different DCA methods decays as a function of the effective depth for families of various lengths. While PlmDCA exhibits the best resilience when the effective depth is lowered, having an adequate accuracy even below *M*_*eff*_ *<* 1000, the autoregressive version of AttentionDCA degrades for *M*_*eff*_ ≤ 2000. Because of this, performances on shallow families such as PF00595, PF13354 or PF00035 are intrinsically worse than those from PlmDCA and the standard version of AttentionDCA.

**Figure 7:**
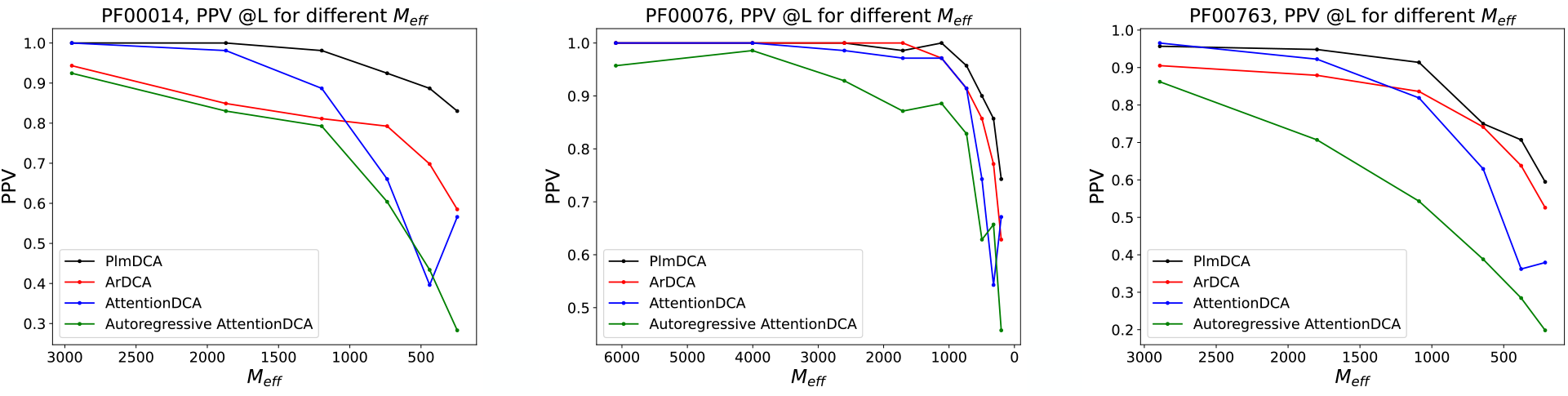
PPV curves @L as the *M*_*eff*_ of the family varies, computed using PlmDCA (FS), ArDCA (ES), AttentionDCA (FS) and Autoregressive AttentionDCA (ES). Families PF00014, PF00076, and PF00763 are shown.

**Figure 8:**
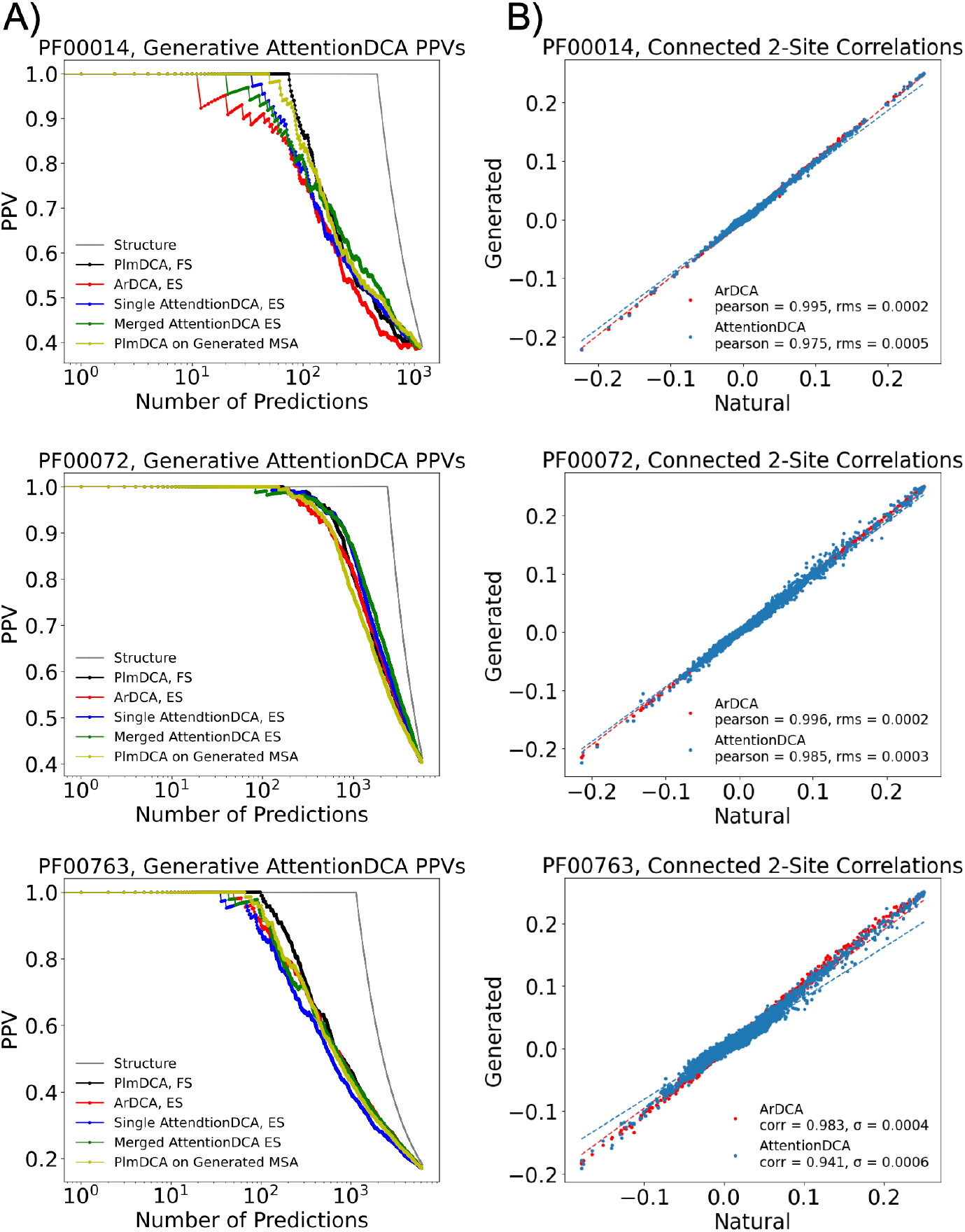
Results from Autoregressive AttentionDCA applied to families PF00014, PF00072 and PF00763. For each family, the panel shows the PPV curves and the connected two-site correlations. The plots on the left compare the PPV curves from PlmDCA (FS, black curve), ArDCA (ES, red curve) with the single- and merged-realizations of AttentionDCA with ES (blue and green curves respectively) and the PPV from the FS of PlmDCA applied to a Generated MSA sampled from Autoregressive AttentionDCA. The gray curve represents the PPV of an ideal model that predicts all and only positive contacts, hence it depicts the prediction of the full structure of the family. The plots on the right show a comparison between the two-site connected correlations of the natural and artificial MSAs generated by Autoregressive AttentionDCA and ArDCA (blue and red dots respectively).

## 5 Discussion

This comprehensive analysis of a single-layer factored self-attention mechanism involved a thorough comparison with the standard pseudo-likelihood approximation found in PlmDCA. Our exploration spanned a diverse dataset, encompassing nine protein families meticulously curated from the Protein Data Bank.

During the past decade, since the development of DCA methods, it became evident the need for parameter reduction through low-rank decomposition [32] or sparse models [33, 34] to avoid overfitting the limited amount of information in an MSA. Remarkably, the primary feature of a factored attention layer compared to a Potts model is the low-rank decomposition that ensures a significant parameter reduction without sacrificing structure prediction accuracy. In particular, the attention mechanism learns structural determinants directly at the level of the attention matrices which can be directly used as a contact score, as shown in Section 4.1.2. Interestingly, as shown in Figure 4 and 5, we found that the contact information is encoded in an effectively sparse organization.

Another important aspect of the factored attention mechanism lies in the differentiation between positional and amino acid epistatic signals from an MSA. As described in Section 3.2, this leads to the possibility of integrating universal amino acid interaction information from different protein families by sharing common sets of parameters in what represents the first example of a multi-family DCA method.

Beyond its primary application in contact prediction, the factored attention layer seamlessly transitions into a fully generative architecture by implementing an autoregressive masking scheme, cf. Section 3.3. Although the initially observed epistatic score contact prediction marginally lagged behind that of similar architecture such as ArDCA [11], our subsequent experimentation involving sequence generation revealed comparable results, as demonstrated in Fig. 8. Ultimately, the observed heterogeneities in the principal component analysis of the generated dataset mirrored the clustering captured by ArDCA, affirming the versatility of our generative factored attention model across various tasks, as shown in Figure 6.

The theoretical and experimental results shown in [15] and [17], along with our in-depth analysis of the potentiality of the factored attention mechanism for protein contact prediction and generation constitute a step forward in breaching the gap between the astonishing results that the attention mechanism has brought in the field of computational biology and the actual understanding we can claim to have of it.

Among the many challenges for future developments, it will be crucial to understand the biological information that can be stored inside value matrices and how this varies when learned from single or multiple protein families. Likewise, it will be interesting to determine whether single attention heads can focus on specific structures or if it is always needed to integrate the information from all heads to produce a meaningful representation of a protein.

## 7 Author contributions statement

FC and AP contributed equally.

## 8 Acknowledgments

The authors acknowledge illuminating discussions with Sergey Ovchinnikov, Martin Weigt, and Francesco Zamponi.

## SUPPLEMENTARY MATERIAL

### A. Summary Statistics

A Multiple Sequence Alignment can be used to extract summary statistics representing the protein family. In particular, single- and pair-wise frequency counts of the amino acids in the alignment are defined as:

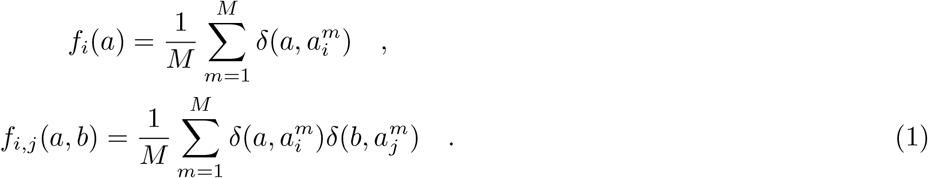

The application of a maximum entropy principle to the frequency counts returns the Potts model at the base of every DCA method. Higher statistics would produce other terms in the Hamiltonian which could not be inferred due to the relatively small size of the datasets at hand.

A good generative model should be able to reproduce the summary statistics and higher statistics derived from it, as in the case of the connected 2-site correlations shown in Figure 8 B) and given by:

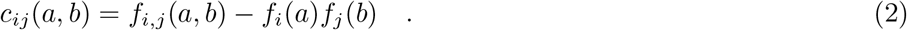

### B. Sequence re-weighting and *M*_*eff*_

Sequences in a natural MSA are not truly independently distributed. Indeed, a certain degree of homology arises due to the phylogenetic relationships between different sequences and it brings about probabilistic biases towards specific sets of sequences. To avoid the over-sampling of similar sequences and the relative under-representation of other sequences, it is necessary to introduce a reweighting scheme. For each sequence **a**^*i*^, a weight is defined as the inverse of the number of sequences similar to it:

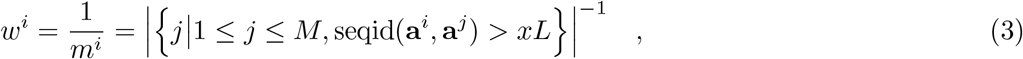

where *x* is a similarity threshold that we set equal to 0.9. These weights enter the computation of the pseudo-likelihood as in:

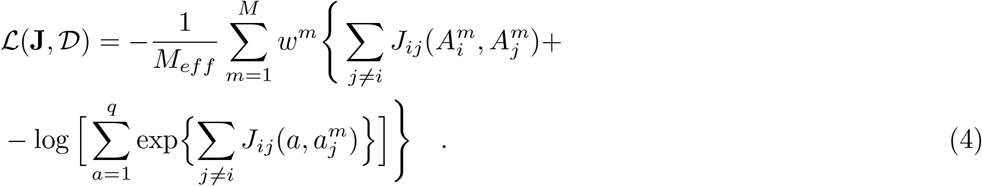

where the effective depth *M*_*eff*_ is then defined as the sum of each weight:

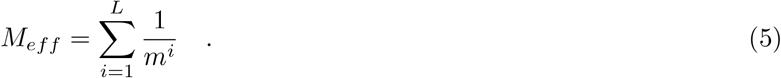

Using this re-weighting scheme, the frequency counts can be written as:

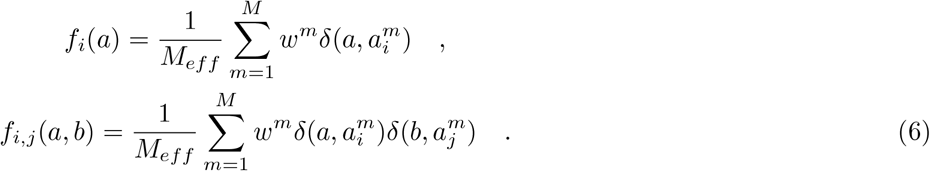

Depending on the effective depth of an MSA, a given DCA method can be more or less efficient, with different methods presenting different precision thresholds.

### C. Average Product Correction

The standard procedure to extract contact information from the interaction tensor of a DCA method is to compute its Frobenious norm to map each *J*_*ij*_(*a, b*) ∈ ℝ_*q×q*_ matrix into a scalar value, useful to define a contact score for positions (*i, j*) as in Equation 7. Empirically, it can be seen that some residues are more prone to establish interactions than others, resulting in a confounding factor for contact prediction. The *average product correction* mitigates this effect by subtracting from the Frobenious norm a null-model contribution for each position pair, so that:

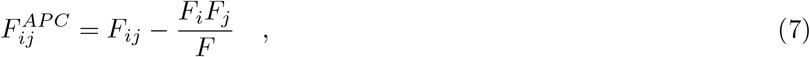

with

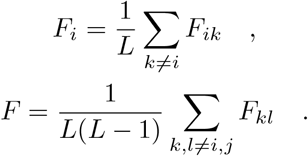

### D. Epistatic Score

As discussed in Sec. II D of the main text, in the generative implementation of the model, the Epistatic Score replaces the interaction tensor when computing the Frobenious norm for direct contact prediction. For a specific pair of amino acids *b*_*i*_, *b*_*j*_ in position (*i, j*), the epistatic score is defined as the difference between the effects of simultaneous mutations on both sites and the sum of the single site mutations when introduced in a given wild-type ***a*** = (*a*_1_, *a*_2_, …, *a*_*L*_):

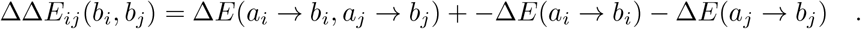

Thus, the ΔΔ*E*_*ij*_(*b*_*i*_, *b*_*j*_) is used instead of the interaction tensor *J*_*ij*_(*a*_*i*_, *a*_*j*_) when computing the Frobenious norm to define the Epistatic Score *ES*_*ij*_. The particular choice of the wild-type sequence can be shown as immaterial for the final contact prediction. In particular, we show that even a randomly generated wild-type would produce the same accuracy as a homolog from the MSA. This signifies that the relevant information is already contained inside the interaction terms and that the computation used for defining the final Epistatic Score measure presents an invariance on the choice of the wild-type. For each Protein Family, figure 1 shows the Epistatic Score PPV@L computed starting from different wild-type sequences, as discussed in Eq. 14 of the main text. In this particular figure, out of two hundred sequences, the first half represents sequences randomly chosen from the corresponding MSA, while the second half represents sequences that have been randomly generated from a uniform distribution. The accuracy of the contact prediction manifestly doesn’t depend on the specific wild-type used in computing the Epistatic Score, whether the sequence actually belongs to the family or is randomly generated.

**FIG. 1.**
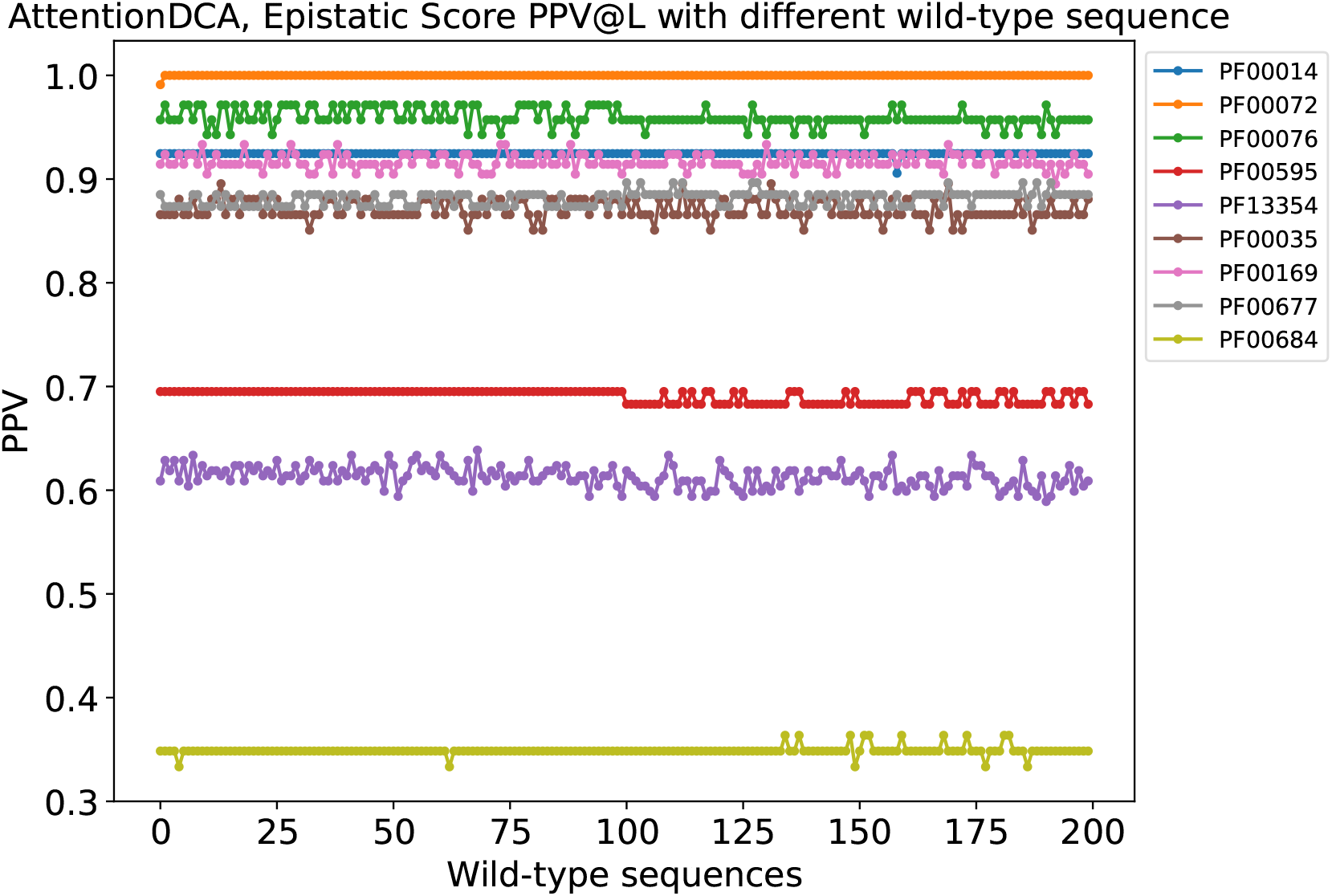
Epistatic Score PPV@L for AttentionDCA using different initial wild-type sequences for the computation of the Epistatic Score. The first half of the wild-type sequences are randomly chosen from the MSA, while the second half sequences are generated from a uniform distribution. Each curve represents a Protein Family according to the legend chart.

### E. Learning Parameters

Table I shows the parameters used for the inference in the standard and autoregressive version of AttentionDCA. If not specified otherwise, the same parameters have been used in both versions. The size of the mini-batches used has been fixed to *b*_*s*_ = 1000 for all families, along with the compression ratio *c*_*r*_ = 0.20 and the number of heads *H* = 128. For each family, the number of epochs and the regularisation have been selected to get an optimal contact prediction.

**TABLE I.**
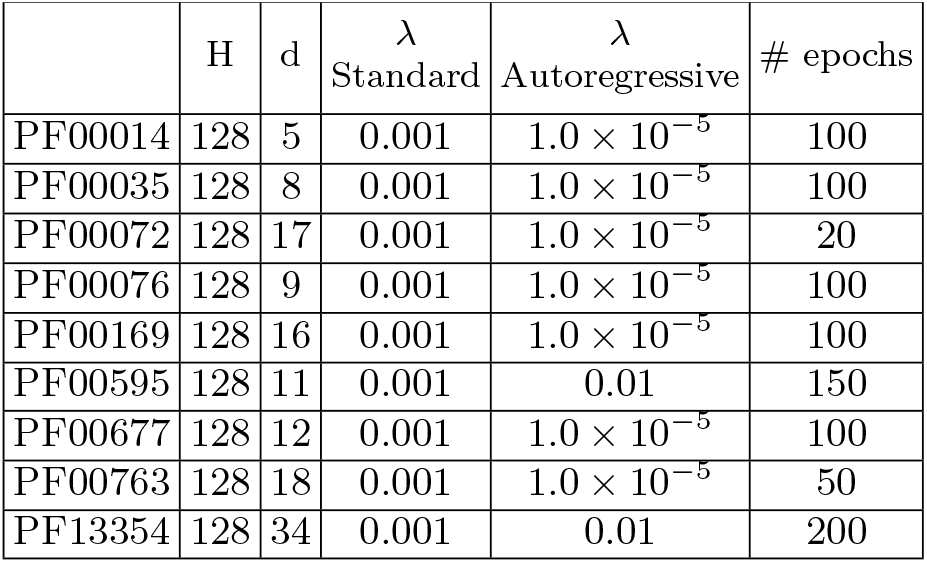
Inference parameters and hyper-parameters of the model for each protein family.

### F. Figures

In the following, we include additional figures showing the results on the complete dataset of protein families available to us. Respectively, we present PPV curves for the non-generative version (cf. Fig. 2 in the main text), contact map from the Frobenious score of the non-generative version (cf. Fig. 3 A)), attention maps (cf. 3 B)), sparse attention maps (cf. Fig. 3 C)), sparse attention PPV curves (cf. Fig. 5), PPV @L for sparse attention at different *k* (cf. Fig. 4), PPV curves for the generative version (cf. Fig. 8 A), 2-site connected correlations (cf. Fig. 8 B)), principal component analysis on ArDCA and PlmDCA (cf. Fig. 6).

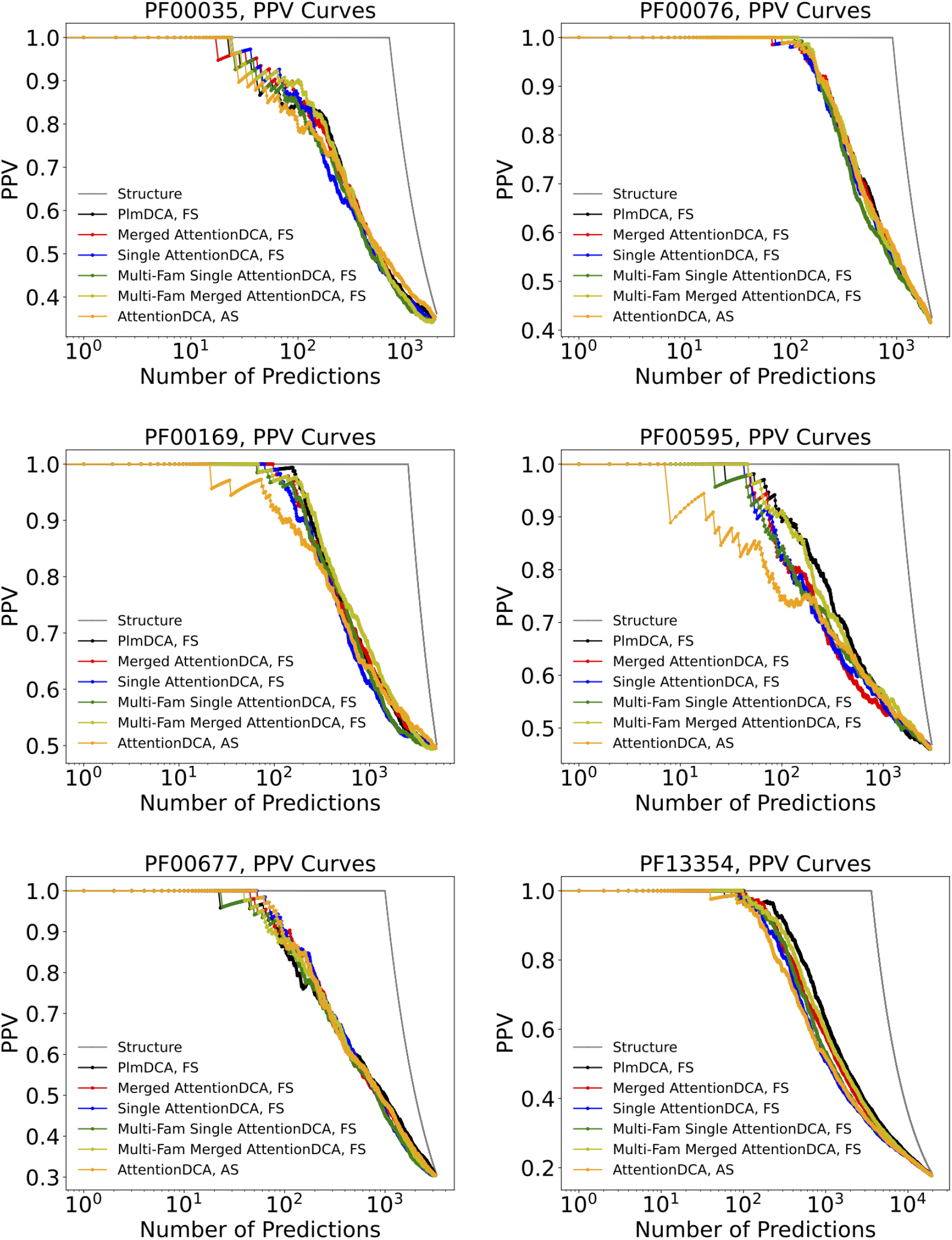

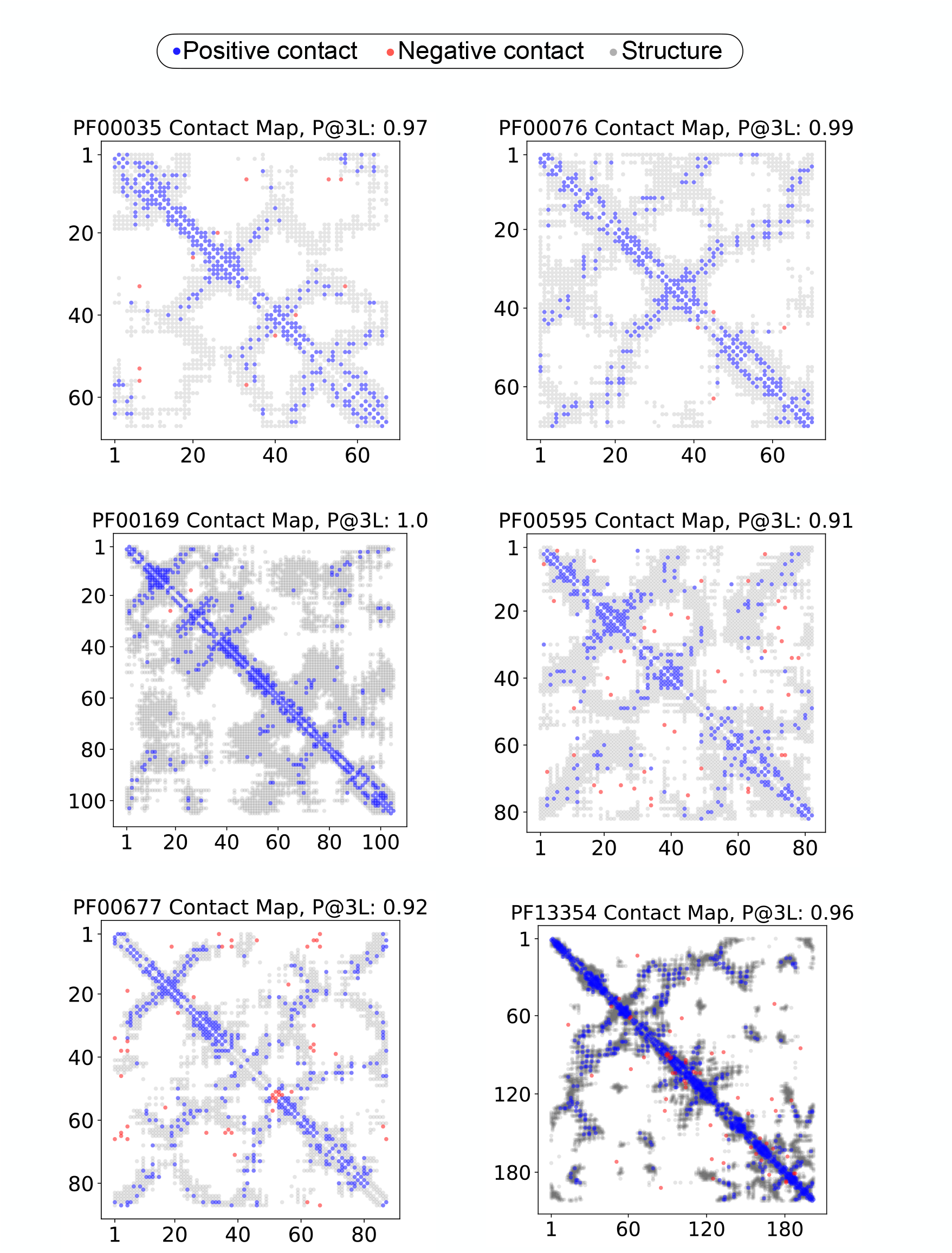

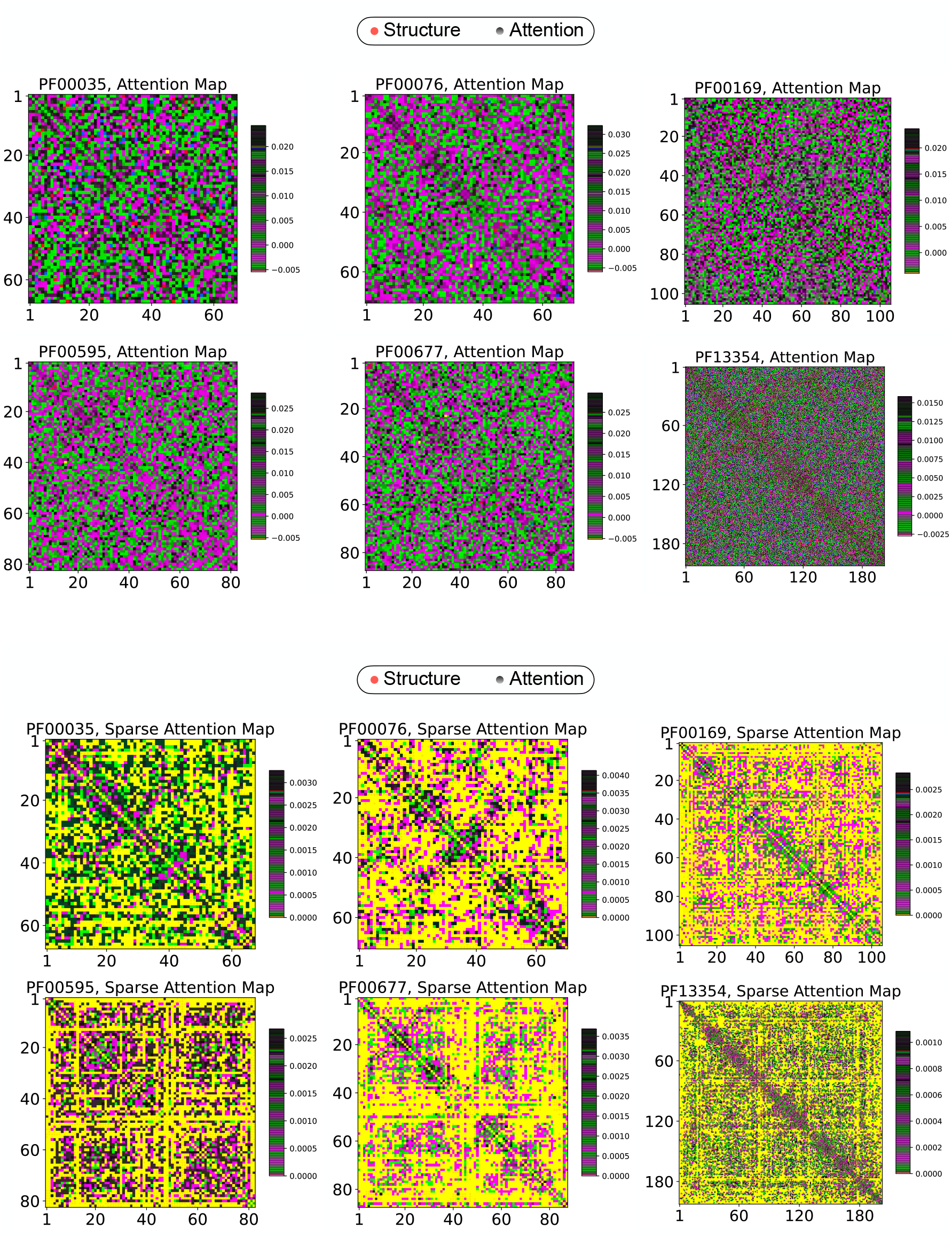

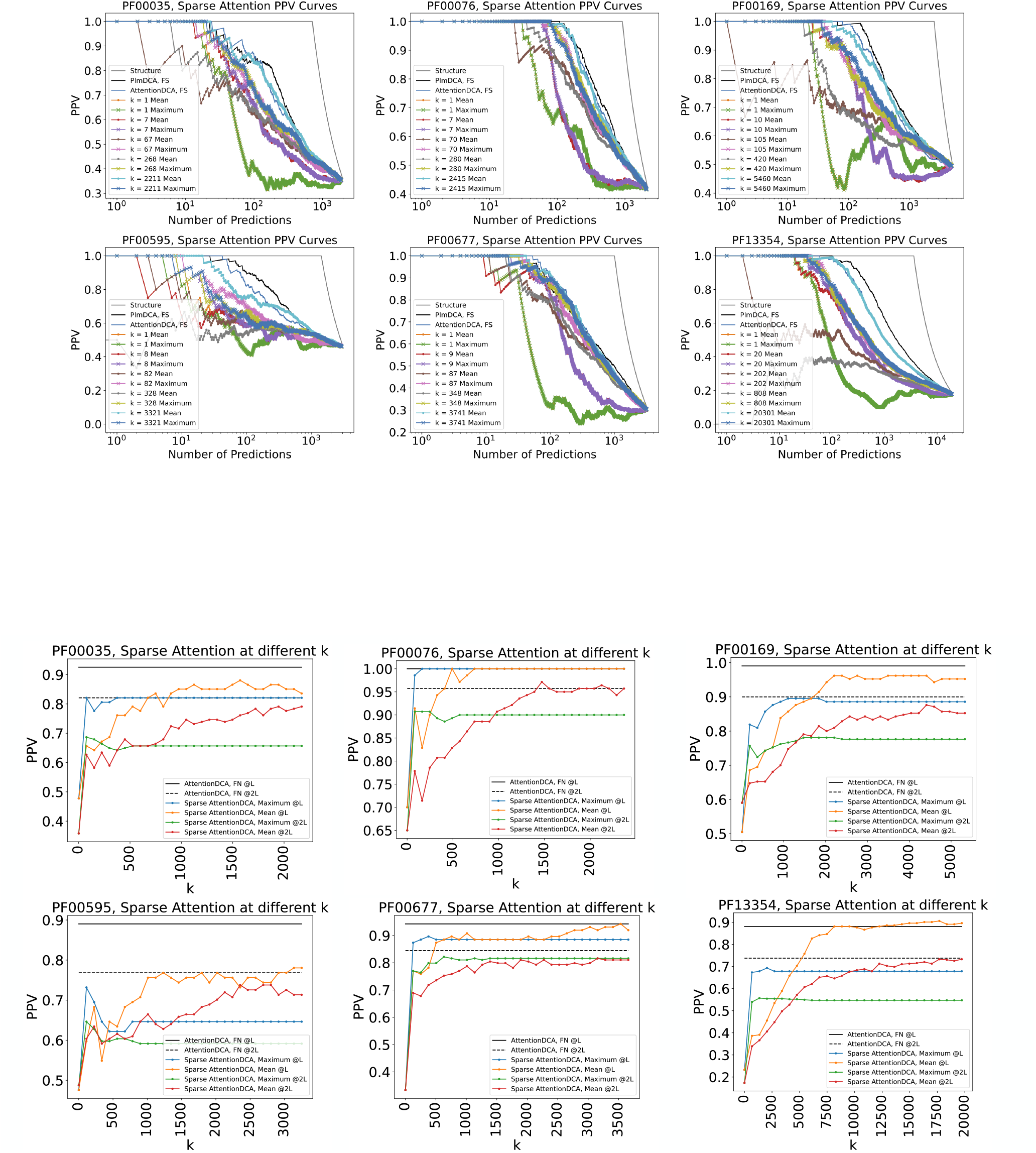

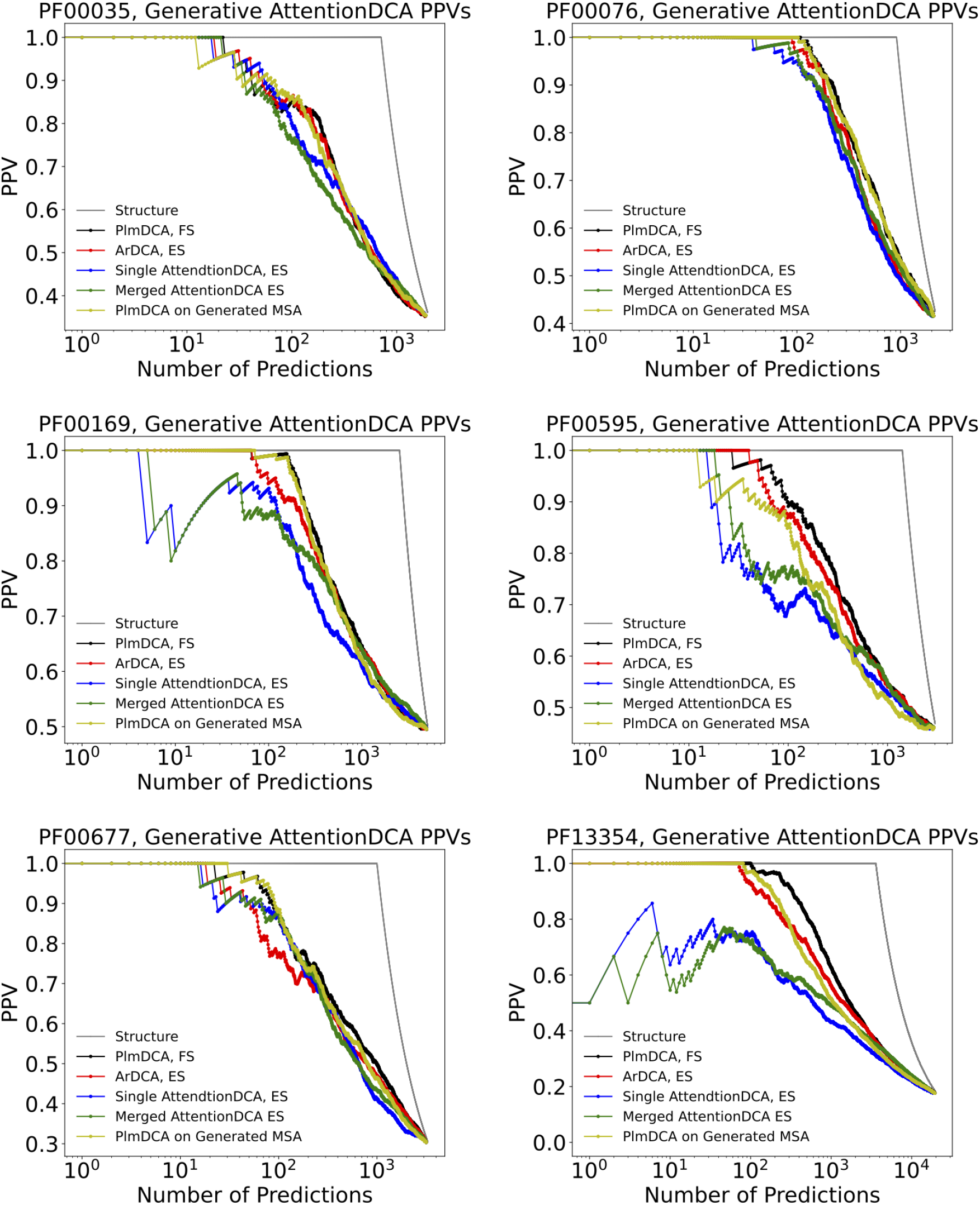

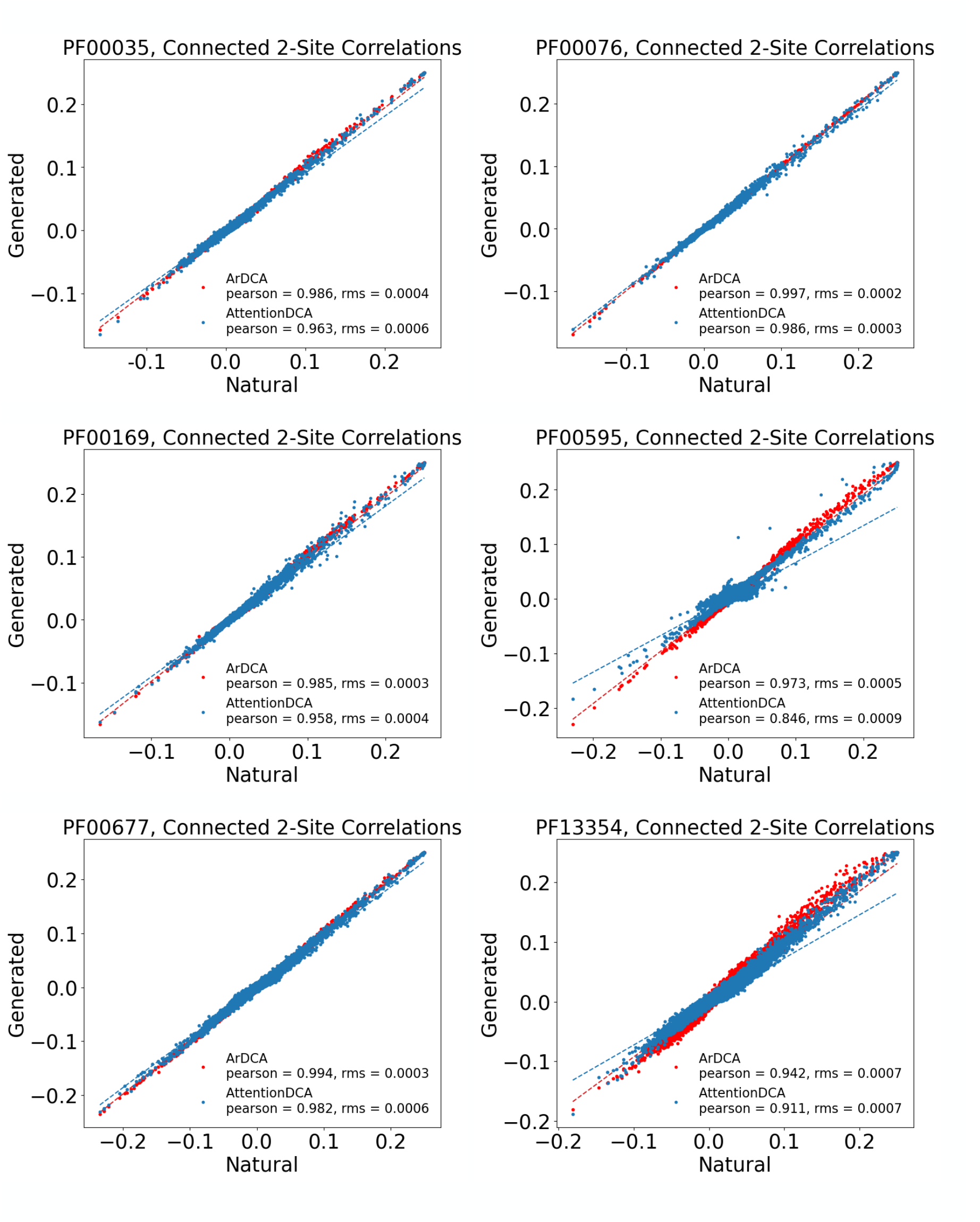

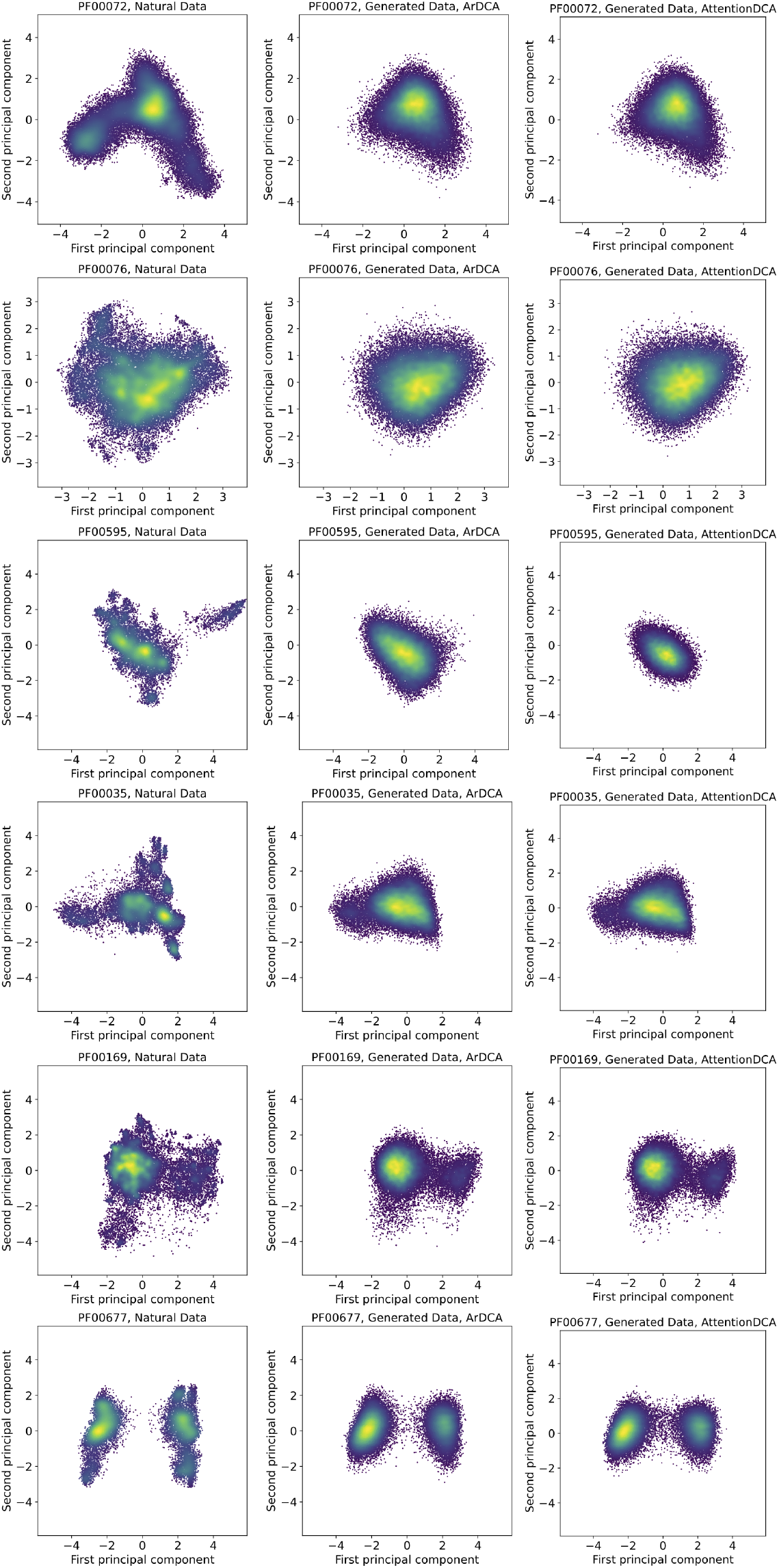

