## supplementary material for "Direct Coupling Analysis and the Attention Mechanism"

### Supplementary Material: Direct Coupling Analysis and the Attention Mechanism

Francesco Caredda[1,*] and Andrea Pagnani[1,2,3,†]
[1] *Politecnico di Torino, Corso Duca degli Abruzzi, 24, I-10129, Torino, Italy.*
[2] *Italian Institute for Genomic Medicine, IRCCS Candiolo, SP-142, I-10060, Candiolo, Italy.*
[3] *INFN, Sezione di Torino, Torino, Via Pietro Giuria, 1 10125 Torino Italy.*
(Dated: September 12, 2024)

#### SUPPLEMENTARY MATERIAL

##### A. Summary Statistics

A Multiple Sequence Alignment can be used to extract summary statistics representing the protein family. In particular, single- and pair-wise frequency counts of the amino acids in the alignment are defined as:

$$f_i(a) = \frac{1}{M}\sum_{m=1}^{M}\delta(a, a_i^m) \quad ,$$
$$f_{i,j}(a,b) = \frac{1}{M}\sum_{m=1}^{M}\delta(a, a_i^m)\delta(b, a_j^m) \quad . \tag{1}$$

The application of a maximum entropy principle to the frequency counts returns the Potts model at the base of every DCA method. Higher statistics would produce other terms in the Hamiltonian which could not be inferred due to the relatively small size of the datasets at hand.

A good generative model should be able to reproduce the summary statistics and higher statistics derived from it, as in the case of the connected 2-site correlations shown in Figure 8 B) and given by:

$$c_{ij}(a,b) = f_{i,j}(a,b) - f_i(a)f_j(b) \quad . \tag{2}$$

##### B. Sequence re-weighting and $M_{eff}$

Sequences in a natural MSA are not truly independently distributed. Indeed, a certain degree of homology arises due to the phylogenetic relationships between different sequences and it brings about probabilistic biases towards specific sets of sequences. To avoid the over-sampling of similar sequences and the relative under-representation of other sequences, it is necessary to introduce a reweighting scheme. For each sequence $\mathbf{a}^i$, a weight is defined as the inverse of the number of sequences similar to it:

$$w^i = \frac{1}{m^i} = \Big|\Big\{ j \Big| 1 \leq j \leq M, \mathrm{seqid}(\mathbf{a}^i, \mathbf{a}^j) > xL \Big\}\Big|^{-1} \quad , \tag{3}$$

where $x$ is a similarity threshold that we set equal to 0.9. These weights enter the computation of the pseudo-likelihood as in:

$$\mathcal{L}(\mathbf{J}, \mathcal{D}) = -\frac{1}{M_{eff}}\sum_{m=1}^{M} w^m \Bigg\{ \sum_{j\neq i} J_{ij}(A_i^m, A_j^m) + \\ - \log\Big[\sum_{a=1}^{q}\exp\Big\{\sum_{j\neq i} J_{ij}(a, a_j^m)\Big\}\Big]\Bigg\} \quad . \tag{4}$$

*
†

where the effective depth $M_{eff}$ is then defined as the sum of each weight:

$$M_{eff} = \sum_{i=1}^{L} \frac{1}{m^i} \quad . \tag{5}$$

Using this re-weighting scheme, the frequency counts can be written as:

$$f_i(a) = \frac{1}{M_{eff}} \sum_{m=1}^{M} w^m \delta(a, a_i^m) \quad ,$$
$$f_{i,j}(a,b) = \frac{1}{M_{eff}} \sum_{m=1}^{M} w^m \delta(a, a_i^m) \delta(b, a_j^m) \quad . \tag{6}$$

Depending on the effective depth of an MSA, a given DCA method can be more or less efficient, with different methods presenting different precision thresholds.

##### C. Average Product Correction

The standard procedure to extract contact information from the interaction tensor of a DCA method is to compute its Frobenious norm to map each $J_{ij}(a,b) \in \mathbb{R}^{q\times q}$ matrix into a scalar value, useful to define a contact score for positions $(i,j)$ as in Equation 7. Empirically, it can be seen that some residues are more prone to establish interactions than others, resulting in a confounding factor for contact prediction. The *average product correction* mitigates this effect by subtracting from the Frobenious norm a null-model contribution for each position pair, so that:

$$F_{ij}^{APC} = F_{ij} - \frac{F_i F_j}{F} \quad , \tag{7}$$

with

$$F_i = \frac{1}{L} \sum_{k \neq i} F_{ik} \quad ,$$
$$F = \frac{1}{L(L-1)} \sum_{k,l \neq i,j} F_{kl} \quad .$$

##### D. Epistatic Score

As discussed in Sec. II D of the main text, in the generative implementation of the model, the Epistatic Score replaces the interaction tensor when computing the Frobenious norm for direct contact prediction. For a specific pair of amino acids $b_i, b_j$ in position $(i,j)$, the epistatic score is defined as the difference between the effects of simultaneous mutations on both sites and the sum of the single site mutations when introduced in a given wild-type $\boldsymbol{a} = (a_1, a_2, \ldots, a_L)$:

$$\Delta\Delta E_{ij}(b_i, b_j) = \Delta E(a_i \to b_i, a_j \to b_j) + -\Delta E(a_i \to b_i) - \Delta E(a_j \to b_j) \quad .$$

Thus, the $\Delta\Delta E_{ij}(b_i,b_j)$ is used instead of the interaction tensor $J_{ij}(a_i,a_j)$ when computing the Frobenious norm to define the Epistatic Score $ES_{ij}$. The particular choice of the wild-type sequence can be shown as immaterial for the final contact prediction. In particular, we show that even a randomly generated wild-type would produce the same accuracy as a homolog from the MSA. This signifies that the relevant information is already contained inside the interaction terms and that the computation used for defining the final Epistatic Score measure presents an invariance on the choice of the wild-type. For each Protein Family, figure 1 shows the Epistatic Score PPV@L computed starting from different wild-type sequences, as discussed in Eq. 14 of the main text. In this particular figure, out of two hundred sequences, the first half represents sequences randomly chosen from the corresponding MSA, while the second half represents sequences that have been randomly generated from a uniform distribution. The accuracy of the contact prediction manifestly doesn't depend on the specific wild-type used in computing the Epistatic Score, whether the sequence actually belongs to the family or is randomly generated.

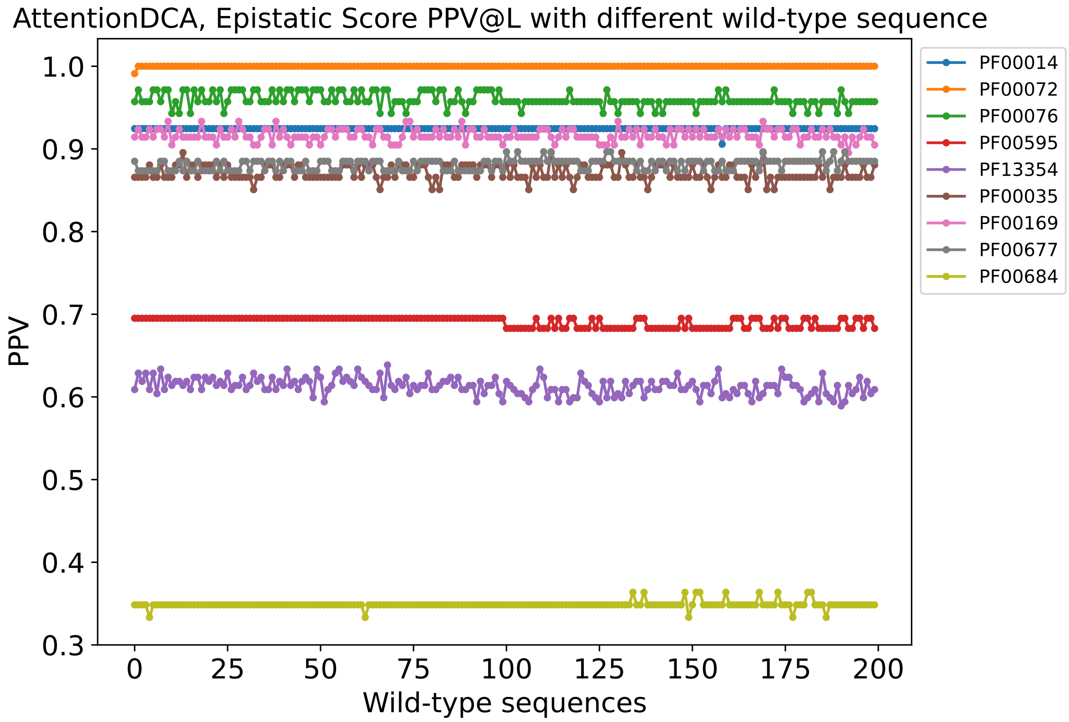


FIG. 1. Epistatic Score PPV@L for AttentionDCA using different initial wild-type sequences for the computation of the Epistatic Score. The first half of the wild-type sequences are randomly chosen from the MSA, while the second half sequences are generated from a uniform distribution. Each curve represents a Protein Family according to the legend chart.

##### E. Learning Parameters

Table I shows the parameters used for the inference in the standard and autoregressive version of AttentionDCA. If not specified otherwise, the same parameters have been used in both versions. The size of the mini-batches used has been fixed to $b_s = 1000$ for all families, along with the compression ratio $c_r = 0.20$ and the number of heads $H = 128$. For each family, the number of epochs and the regularisation have been selected to get an optimal contact prediction.

TABLE I. Inference parameters and hyper-parameters of the model for each protein family.

| | H | d | $\lambda$ Standard | $\lambda$ Autoregressive | # epochs |
|---|---|---|---|---|---|
| PF00014 | 128 | 5 | 0.001 | $1.0 \times 10^{-5}$ | 100 |
| PF00035 | 128 | 8 | 0.001 | $1.0 \times 10^{-5}$ | 100 |
| PF00072 | 128 | 17 | 0.001 | $1.0 \times 10^{-5}$ | 20 |
| PF00076 | 128 | 9 | 0.001 | $1.0 \times 10^{-5}$ | 100 |
| PF00169 | 128 | 16 | 0.001 | $1.0 \times 10^{-5}$ | 100 |
| PF00595 | 128 | 11 | 0.001 | 0.01 | 150 |
| PF00677 | 128 | 12 | 0.001 | $1.0 \times 10^{-5}$ | 100 |
| PF00763 | 128 | 18 | 0.001 | $1.0 \times 10^{-5}$ | 50 |
| PF13354 | 128 | 34 | 0.001 | 0.01 | 200 |

##### F. Figures

In the following, we include additional figures showing the results on the complete dataset of protein families available to us. Respectively, we present PPV curves for the non-generative version (cf. Fig. 2 in the main text), contact map from the Frobenious score of the non-generative version (cf. Fig. 3 A)), attention maps (cf. 3 B)), sparse attention maps (cf. Fig. 3 C)), sparse attention PPV curves (cf. Fig. 5), PPV @L for sparse attention at different $k$ (cf. Fig. 4), PPV curves for the generative version (cf. Fig. 8 A), 2-site connected correlations (cf. Fig. 8 B)), principal component analysis on ArDCA and PlmDCA (cf. Fig. 6).

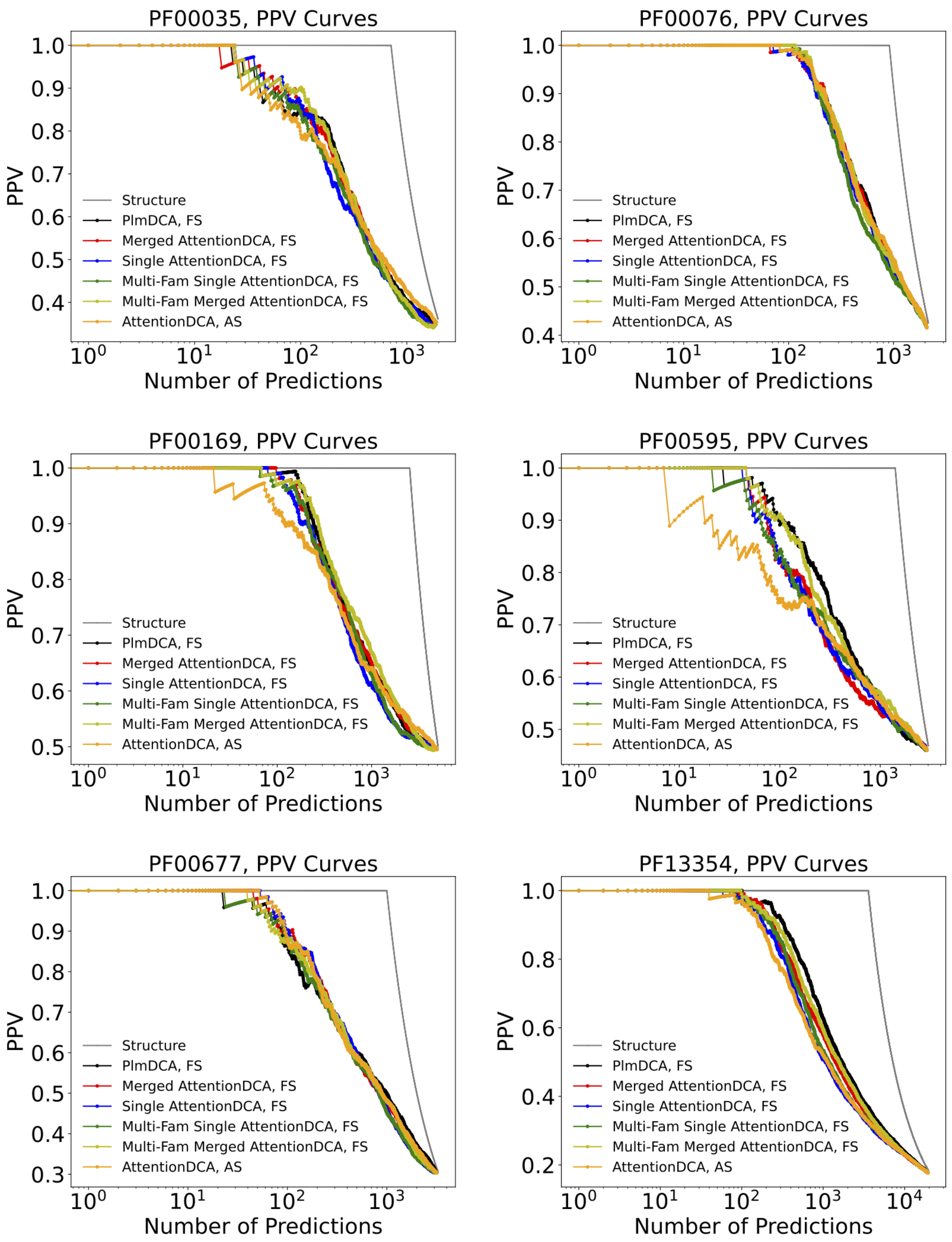

PF00035, PPV Curves
PF00076, PPV Curves
PF00169, PPV Curves
PF00595, PPV Curves
PF00677, PPV Curves
PF13354, PPV Curves
PPV
Number of Predictions
Structure
PlmDCA, FS
Merged AttentionDCA, FS
Single AttentionDCA, FS
Multi-Fam Single AttentionDCA, FS
Multi-Fam Merged AttentionDCA, FS
AttentionDCA, AS

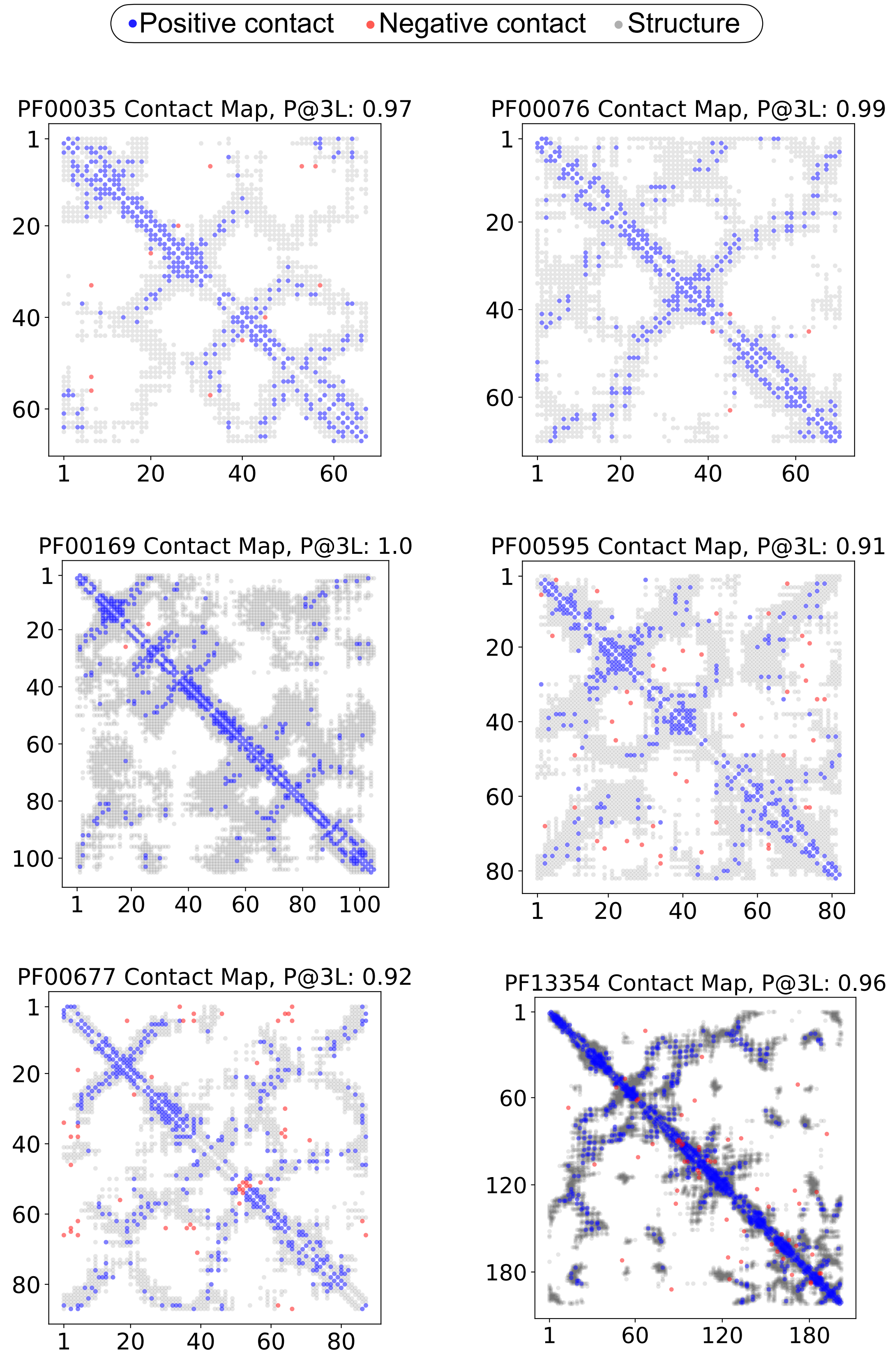
Positive contact
Negative contact
Structure
PF00035 Contact Map, P@3L: 0.97
PF00076 Contact Map, P@3L: 0.99
PF00169 Contact Map, P@3L: 1.0
PF00595 Contact Map, P@3L: 0.91
PF00677 Contact Map, P@3L: 0.92
PF13354 Contact Map, P@3L: 0.96
1
20
40
60
80
100
120
180

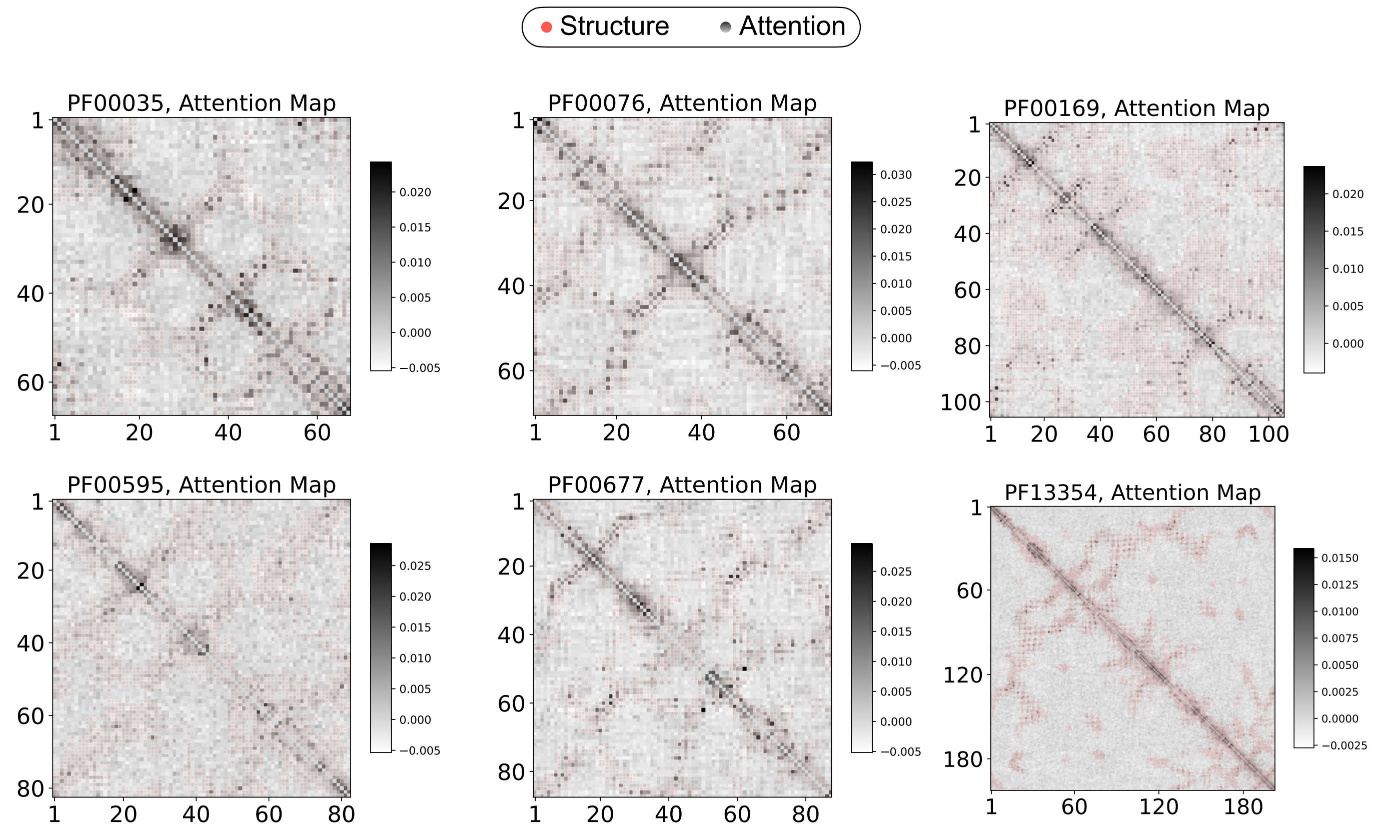
Structure
Attention
PF00035, Attention Map
PF00076, Attention Map
PF00169, Attention Map
PF00595, Attention Map
PF00677, Attention Map
PF13354, Attention Map

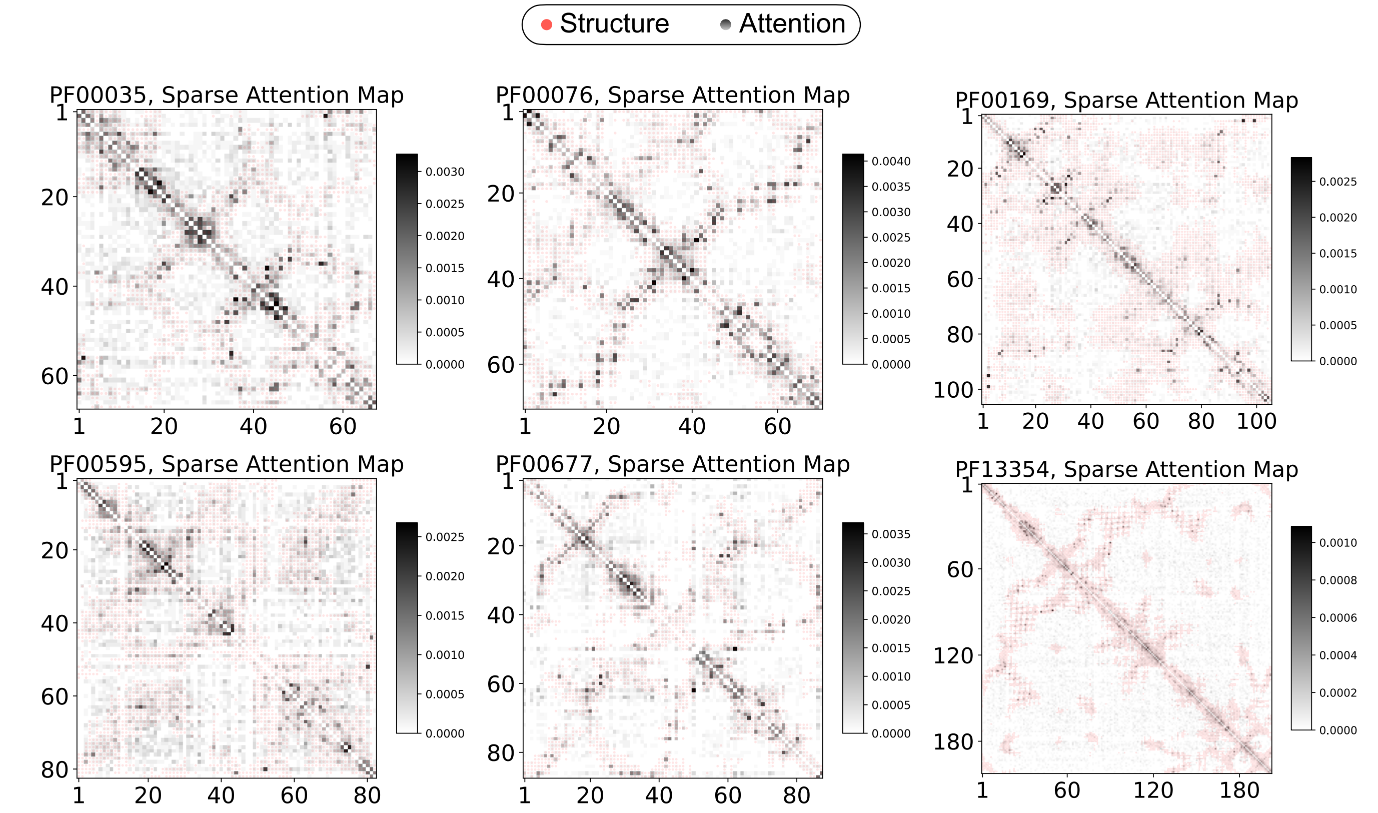
Structure
Attention
PF00035, Sparse Attention Map
PF00076, Sparse Attention Map
PF00169, Sparse Attention Map
PF00595, Sparse Attention Map
PF00677, Sparse Attention Map
PF13354, Sparse Attention Map

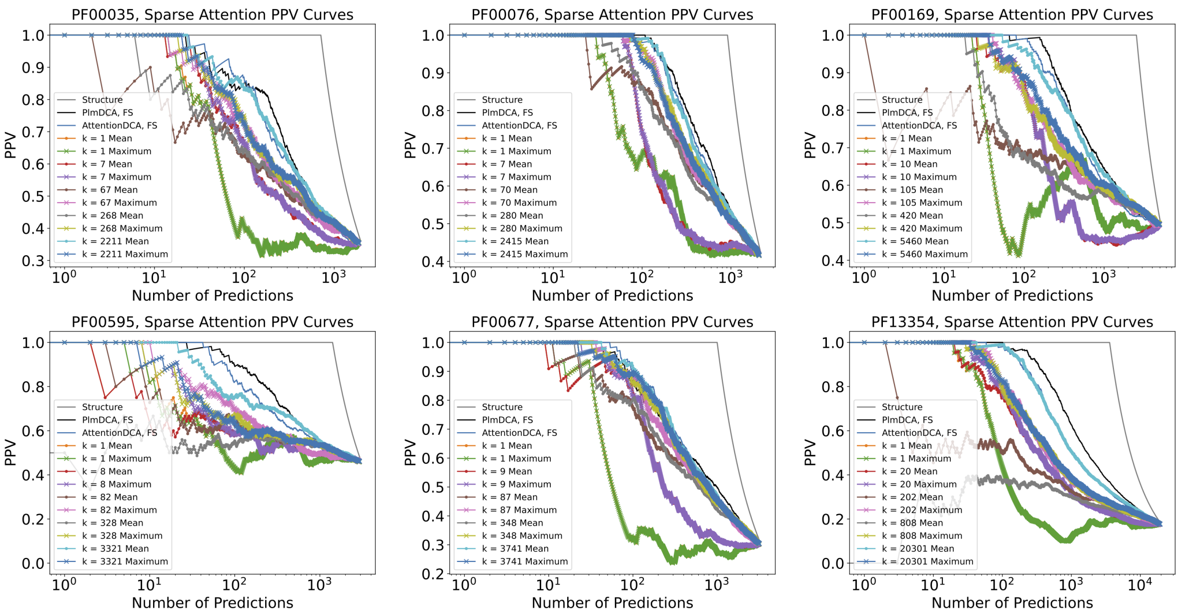

PF00035, Sparse Attention PPV Curves
PF00076, Sparse Attention PPV Curves
PF00169, Sparse Attention PPV Curves
PF00595, Sparse Attention PPV Curves
PF00677, Sparse Attention PPV Curves
PF13354, Sparse Attention PPV Curves
PPV
Number of Predictions
Structure
PlmDCA, FS
AttentionDCA, FS
k = 1 Mean
k = 1 Maximum
k = 7 Mean
k = 7 Maximum
k = 67 Mean
k = 67 Maximum
k = 268 Mean
k = 268 Maximum
k = 2211 Mean
k = 2211 Maximum
k = 70 Mean
k = 70 Maximum
k = 280 Mean
k = 280 Maximum
k = 2415 Mean
k = 2415 Maximum
k = 10 Mean
k = 10 Maximum
k = 105 Mean
k = 105 Maximum
k = 420 Mean
k = 420 Maximum
k = 5460 Mean
k = 5460 Maximum
k = 8 Mean
k = 8 Maximum
k = 82 Mean
k = 82 Maximum
k = 328 Mean
k = 328 Maximum
k = 3321 Mean
k = 3321 Maximum
k = 9 Mean
k = 9 Maximum
k = 87 Mean
k = 87 Maximum
k = 348 Mean
k = 348 Maximum
k = 3741 Mean
k = 3741 Maximum
k = 20 Mean
k = 20 Maximum
k = 202 Mean
k = 202 Maximum
k = 808 Mean
k = 808 Maximum
k = 20301 Mean
k = 20301 Maximum


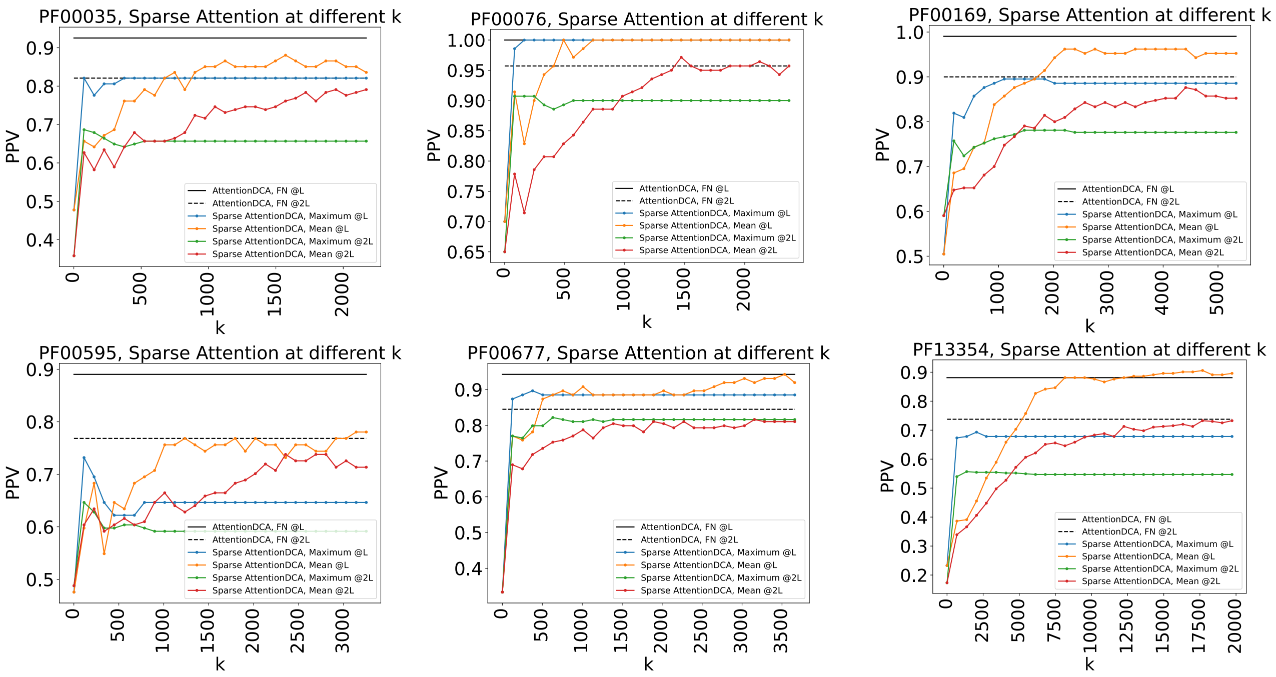

PF00035, Sparse Attention at different k
PF00076, Sparse Attention at different k
PF00169, Sparse Attention at different k
PF00595, Sparse Attention at different k
PF00677, Sparse Attention at different k
PF13354, Sparse Attention at different k
PPV
k
AttentionDCA, FN @L
AttentionDCA, FN @2L
Sparse AttentionDCA, Maximum @L
Sparse AttentionDCA, Mean @L
Sparse AttentionDCA, Maximum @2L
Sparse AttentionDCA, Mean @2L

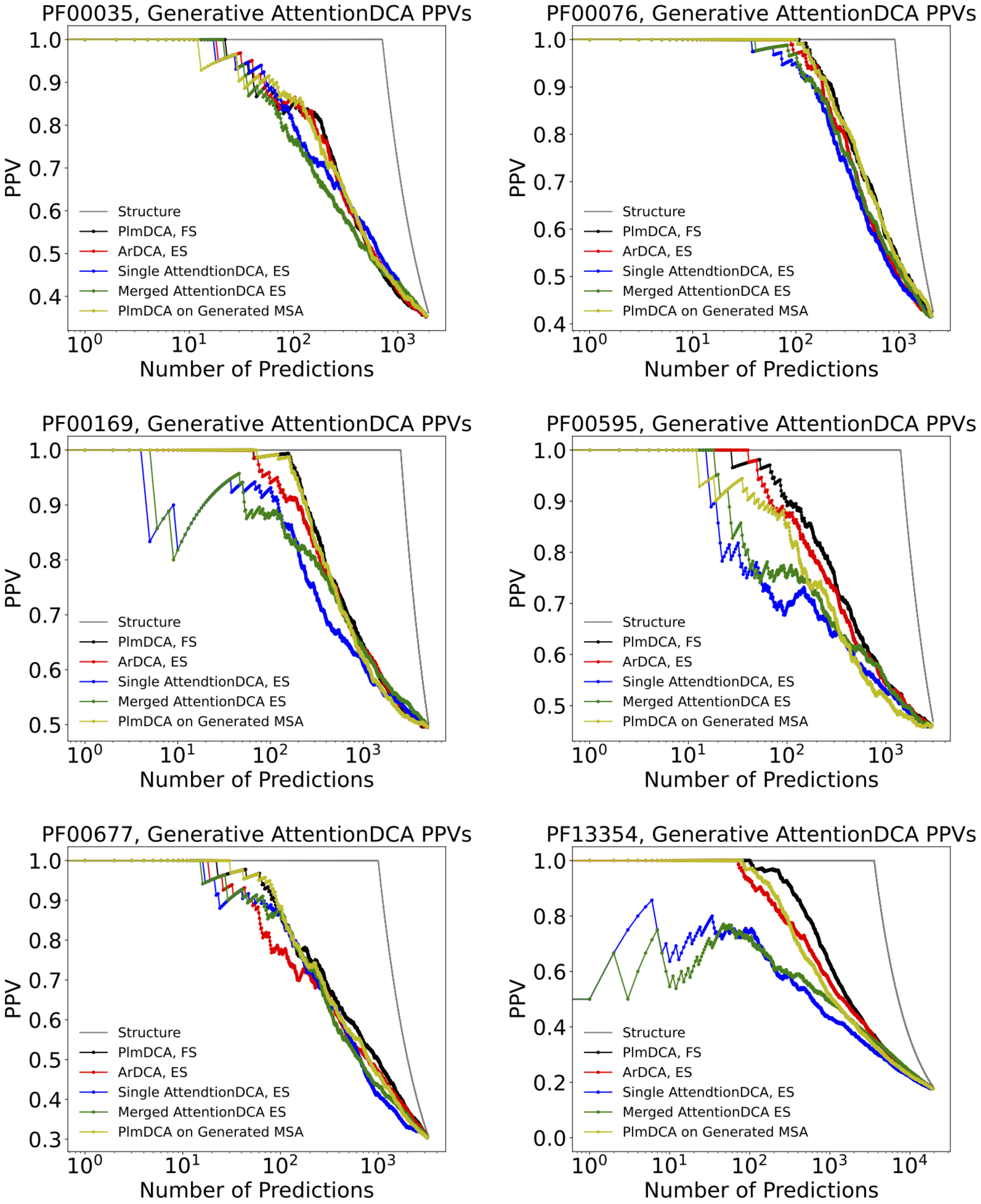

PF00035, Generative AttentionDCA PPVs
PF00076, Generative AttentionDCA PPVs
PF00169, Generative AttentionDCA PPVs
PF00595, Generative AttentionDCA PPVs
PF00677, Generative AttentionDCA PPVs
PF13354, Generative AttentionDCA PPVs
PPV
Number of Predictions
Structure
PlmDCA, FS
ArDCA, ES
Single AttendtionDCA, ES
Merged AttentionDCA ES
PlmDCA on Generated MSA

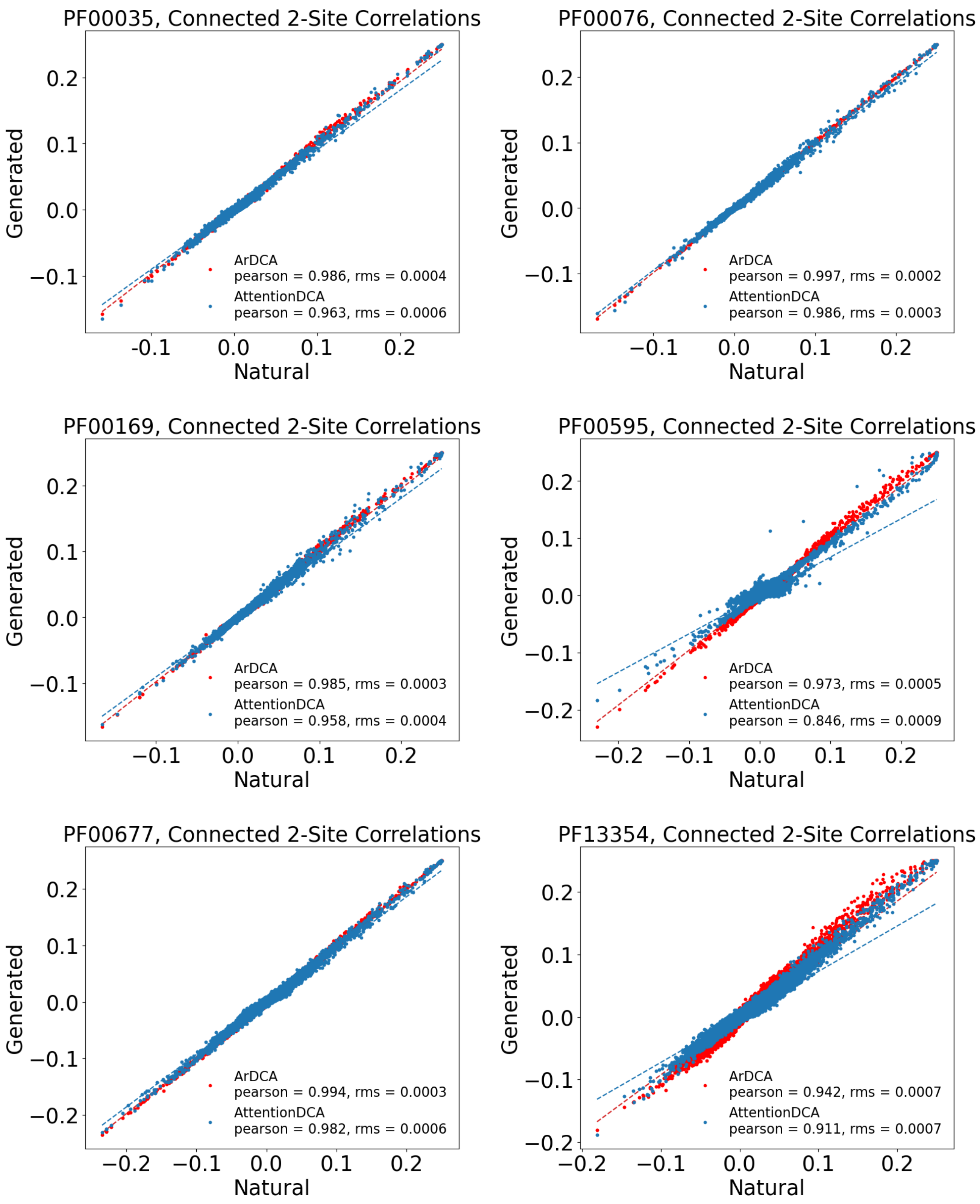
PF00035, Connected 2-Site Correlations
Generated
Natural
ArDCA
pearson = 0.986, rms = 0.0004
AttentionDCA
pearson = 0.963, rms = 0.0006
PF00076, Connected 2-Site Correlations
Generated
Natural
ArDCA
pearson = 0.997, rms = 0.0002
AttentionDCA
pearson = 0.986, rms = 0.0003
PF00169, Connected 2-Site Correlations
Generated
Natural
ArDCA
pearson = 0.985, rms = 0.0003
AttentionDCA
pearson = 0.958, rms = 0.0004
PF00595, Connected 2-Site Correlations
Generated
Natural
ArDCA
pearson = 0.973, rms = 0.0005
AttentionDCA
pearson = 0.846, rms = 0.0009
PF00677, Connected 2-Site Correlations
Generated
Natural
ArDCA
pearson = 0.994, rms = 0.0003
AttentionDCA
pearson = 0.982, rms = 0.0006
PF13354, Connected 2-Site Correlations
Generated
Natural
ArDCA
pearson = 0.942, rms = 0.0007
AttentionDCA
pearson = 0.911, rms = 0.0007

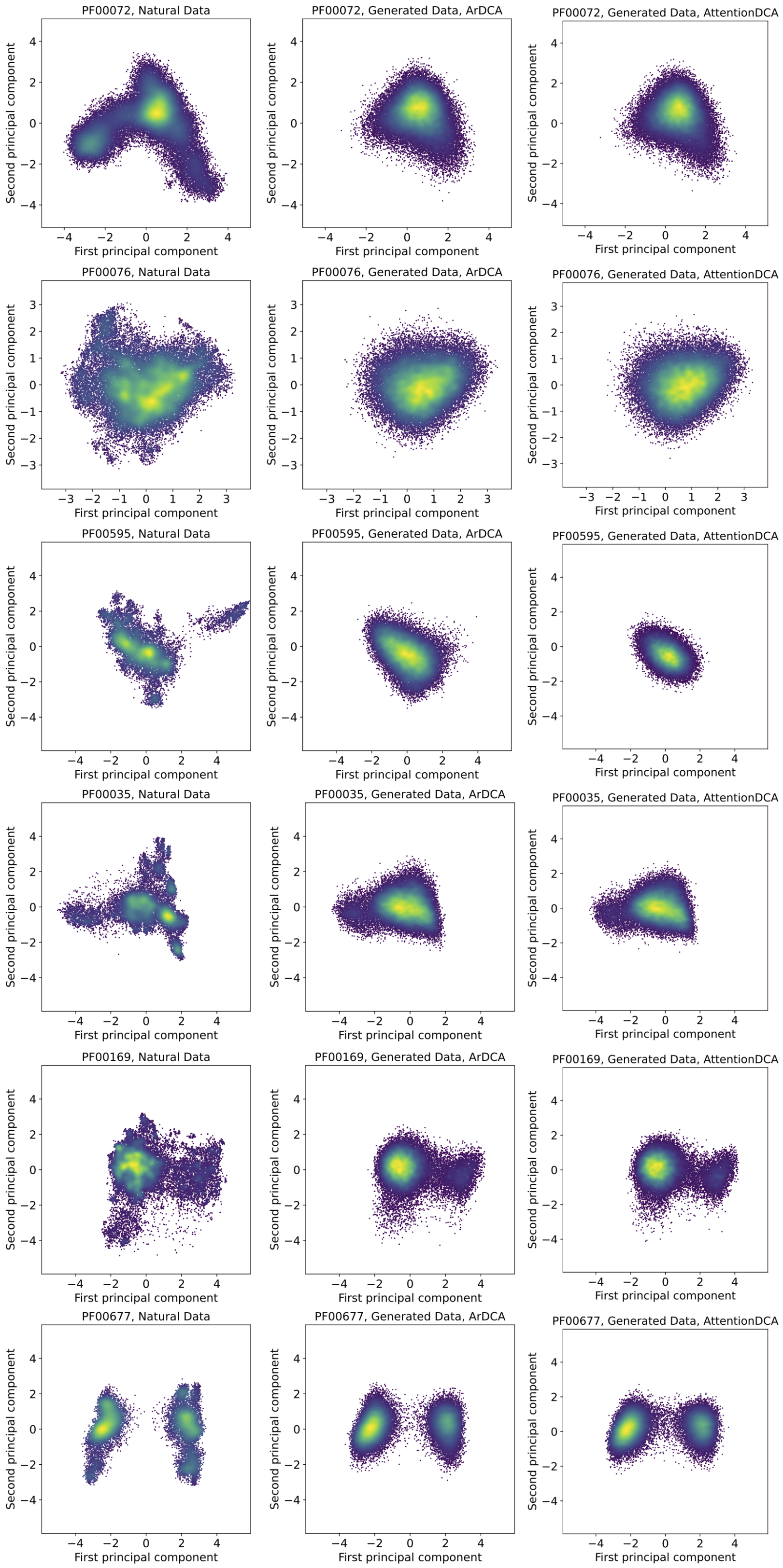
PF00072, Natural Data
PF00072, Generated Data, ArDCA
PF00072, Generated Data, AttentionDCA
PF00076, Natural Data
PF00076, Generated Data, ArDCA
PF00076, Generated Data, AttentionDCA
PF00595, Natural Data
PF00595, Generated Data, ArDCA
PF00595, Generated Data, AttentionDCA
PF00035, Natural Data
PF00035, Generated Data, ArDCA
PF00035, Generated Data, AttentionDCA
PF00169, Natural Data
PF00169, Generated Data, ArDCA
PF00169, Generated Data, AttentionDCA
PF00677, Natural Data
PF00677, Generated Data, ArDCA
PF00677, Generated Data, AttentionDCA
First principal component
Second principal component
